# Expressed Exome Capture Sequencing (EecSeq): a method for cost-effective exome sequencing for all organisms with or without genomic resources

**DOI:** 10.1101/223735

**Authors:** Jonathan B. Puritz, Katie E Lotterhos

## Abstract

Exome capture is an effective tool for surveying the genome for loci under selection. However, traditional methods require annotated genomic resources. Here, we present a method for creating cDNA probes from expressed mRNA, which are then used to enrich and capture genomic DNA for exon regions. This approach, called “EecSeq”, eliminates the need for costly probe design and synthesis. We tested EecSeq in the eastern oyster, *Crassostrea virginica*, using a controlled exposure experiment. Four adult oysters were heat shocked at 36° C for 1 hour along with four control oysters kept at 14° C. Stranded mRNA libraries were prepared for two individuals from each treatment and pooled. Half of the combined library was used for probe synthesis and half was sequenced to evaluate capture efficiency. Genomic DNA was extracted from all individuals, enriched via captured probes, and sequenced directly. We found that EecSeq had an average capture sensitivity of 86.8% across all known exons and had over 99.4% sensitivity for exons with detectable levels of expression in the mRNA library. For all mapped reads, over 47.9% mapped to exons and 37.0% mapped to expressed targets, which is similar to previously published exon capture studies. EecSeq displayed relatively even coverage within exons (i.e. minor “edge effects”) and even coverage across exon GC content. We discovered 5,951 SNPs with a minimum average coverage of 80X, with 3,508 SNPs appearing in exonic regions. We show that EecSeq provides comparable, if not superior, specificity and capture efficiency compared to costly, traditional methods.

## Introduction

The invention of next-generation sequencing has made it possible to obtain massive amounts of sequence data. These data have given insight into classical problems in evolutionary biology, including the repeatability of evolution (e.g., Jones *et al.* 2012), the degree of convergent evolution across distant taxa (e.g., Yeaman *et al.* 2016), and whether selection is driving changes in existing genetic variation or new mutations (e.g., Reid *et al.* 2016). Despite this rapid progress, it is still cost prohibitive to sequence dozens or hundreds of full genomes. This limits our ability to study the genomic basis of local adaptation, which requires large sample sizes for statistical power (De Mita *et al.* 2013; Lotterhos & Whitlock 2015; Hoban *et al.* 2016). This leads to an inherent trade-off between sample size and genomic coverage, leading investigators to make decisions about whether to sequence more individuals (for higher power and precision) versus more of the genome (for making more accurate statements about the genetic basis of adaptation).

Reduced representation library preparation methods offer various kinds of random or targeted genome reduction, but the available approaches have contrasting advantages and limitations. RADseq uses restriction enzymes to randomly sample the genome and is appropriate for linkage mapping and studying neutral processes like gene flow and drift (Puritz *et al.* 2014), but the data can be limited for understanding the genetic basis of adaptation (Lowry *et al.* 2016, 2017; Catchen *et al.* 2017; McKinney *et al.* 2017). To focus on coding regions, some investigators have used RNAseq (De Wit *et al.* 2015); however, only about a dozen individuals can be sequenced per lane because of log-fold differences in transcript abundance among loci. Additionally, allele-specific expression limits the confidence in genotypes derived from RNAseq data (Pastinen 2010), especially in pooled samples. One increasingly popular option for increasing precision with larger samples while still maintaining coverage of the entire genome is Pool-seq, which sequences every individual to very low (1x) coverage and uses the data to calculate allele frequency of the sample within the pool (Buerkle & Gompert 2013; Schlötterer *et al.* 2014; Therkildsen & Palumbi 2017). Pool-seq is limited to only estimating allele frequency within pools, which is a disadvantage because this data cannot be used to understand the fitness of heterozygotes and some types of statistical analyses would be impossible to perform, such as haplotype-based analyses (e.g. Fariello *et al.* 2013).

To overcome some of these limitations, many investigators have used capture approaches with biotinylated probes (Jones & Good 2016). Capture approaches have the advantage of enriching the data for sequences of interest - allowing for individual-level data and a large number of individuals to be sequenced - but require the investigator to have genomic resources for probe design and then to purchase the probes from a company. For non-model species, the development of these resources takes time and a significant amount of bioinformatics expertise. In addition, for a population-level genomic study with 100s of individuals, probes may cost several tens of thousands of dollars, depending on how much sequence is captured. Overall, what is needed is a cost-effective approach to subsample genomes for coding regions, without previously developed genomic resources. Such an approach would allow for the assessment of rapid adaptation to environmental disasters such as Deepwater Horizon Oil Spill (Lee *et al.* 2017).

Here, we present a novel, cost-effective method of exome capture that synthesizes probes in-situ from expressed mRNA sequences. Expressed Exome Capture Sequencing (EecSeq) builds upon existing approaches for in-situ probe synthesis that rely on restriction enzymes to sample the genome or exome (Suchan *et al.* 2016; Schmid *et al.* 2017). To improve capture efficiency, we developed a novel library preparation procedure that uses standardized procedures to synthesize cDNA from expressed RNA (without template reduction via restriction digest) and then create biotinylated probes from cDNA (see Figure 1 for a conceptual diagram). The EecSeq design includes custom RNA library adapters that offer several major advantages. The custom adapters are fully compatible with duplex-specific nuclease normalization, which is included in the protocol in order to reduce log fold differences in expression - resulting in more even coverage across high-and low-expressed transcripts. The custom adapters also allow for probe sequencing - before normalization if differential expression data is desired, or after normalization if probe abundance data is desired. Moreover, the adapters are easily removed with a single enzymatic treatment before biotinylation, preventing any interference during hybridization.

**Figure 1.**
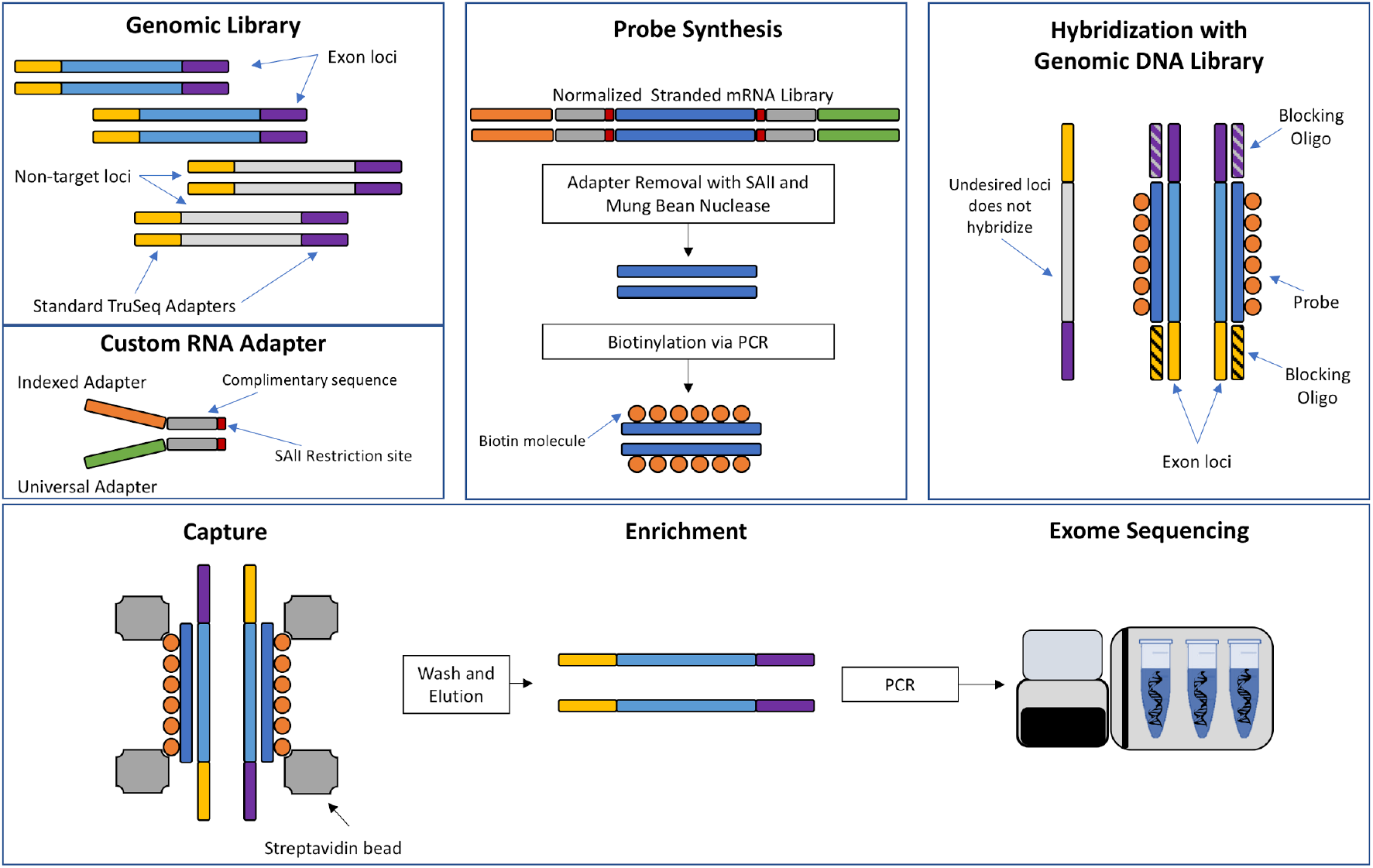
Conceptual Diagram of Expressed Exome Capture Sequencing. Upper left panel: The shotgun genomic DNA library that will be captured with probes. Middle left panel: EecSeq relies on custom RNA adapters that contains a SAlI restriction site. Middle upper panel: The adapters are incorporated into a mRNA library preparation that is normalized with duplex-specific nuclease. Adapters are then removed with a SA1I restriction digest, cDNA probes are subsequently blunted with mung bean nuclease, and biotinylated via a PCR reaction. Upper right panel: The probes are then hybridized to the shotgun genomic library with TruSeq style adapters. Exon loci bind to the cDNA probes. Lower panel: Hybridized exon loci and probes are then captured with magnetic Streptavidin beads. The captured exome fragments are washed several times, eluted, enriched with PCR, and then sequenced.

Our approach is cost-effective and does not require any prior genomic resources, making it a good choice for studies seeking to understand adaptation in exomes. The approach, however, is limited in the sense that the probes are designed from expressed RNA, and so investigators should be careful to choose which tissues and life stages would be relevant. Here, we show proof-of-concept of the approach in the eastern oyster *(Crassostrea virginica)*, and find that the performance of the approach is comparable, if not superior, to the performance of published exome capture datasets where probes were designed from sequence data and purchased from a company.

## Methods

### Experimental overview

Expressed exome capture sequencing (EecSeq) is designed with two specific goals: 1) to eliminate the need for expensive exome capture probe design and synthesis and 2) to focus exon enrichment of genes that are being expressed relevant to tissue(s) and condition(s) of interest. To illustrate this conceptually, we exposed adult oysters to a stressor (extreme heat) that would generate a predictable gene and protein expression profile (expression of heat shock proteins). Having a predictable coverage profile in the probes allowed us to evaluate whether the genomic DNA in these exons were captured by the probes. Note, however, that this experiment is not specifically part of the EecSeq method and that the investigator can choose appropriate tissue(s) and condition(s) of interest. The steps to probe synthesis and capture are visualized in Figure 1.

### Heat shock exposure, tissue collection, and nucleic acid extraction

Eight adult *Crassostrea virginica* individuals were collected and acclimated to a flow-through seawater system for 24 hours. After acclimation, individuals were randomly assigned to two treatments, control and heat-shock (HS). HS individuals were placed a small aquaria filled with 36°C filtered seawater for one hour while control individuals were kept in an identical aquarium filled with 14°C (ambient) filtered seawater. Immediately after the exposure period, all individuals were shucked and mantle tissue was extracted and frozen in liquid nitrogen in duplicate. DNA was extracted using the DNeasy kit (Qiagen) and RNA was extracted using TRI Reagent Solution (Applied Biosystems) using included, standard protocols. DNA was visualized on an agarose gel and quantified using the Qubit DNA Broad Range kit (Invitrogen). RNA was visualized on an Agilent BioAnalyzer using the RNA 6000 Nano kit, and was quantified using the Qubit High Sensitivity Assay Kit (Invitrogen).

### Expressed Exome Capture Sequencing

A complete and updated EecSeq protocol can be found at (https://github.com/jpuritz/EecSeq). *RNA Adapters-* Custom RNA adapters were used in this protocol. The RNA adapters were similar to the Illumina TruSeq design, but include the SAlI restriction site at the 3’ end of the “Universal adapter” and at 5’ end of the “Indexed adapter.” The presence of this restriction site allows the Illumina sequence to be removed before hybridization to prevent interference. Note that the adapters used in this study had an erroneous deletion of a Thymine in position 58 of “Universal_SAI1_Adapter” and in position 8 of all four indexed adapters (the corrected versions are shown in Table 1, and erroneous version used in this study are shown in Supplemental Table 1). Adapters were annealed in equal parts in a solution of Tris-HCl (pH 8.0), NaCl, and EDTA, heated to 97.5°C for 2.5 minutes, and then cooled at a rate of 3°C per minute until the solution reaches a temperature of 21°C.

**Table 1.**
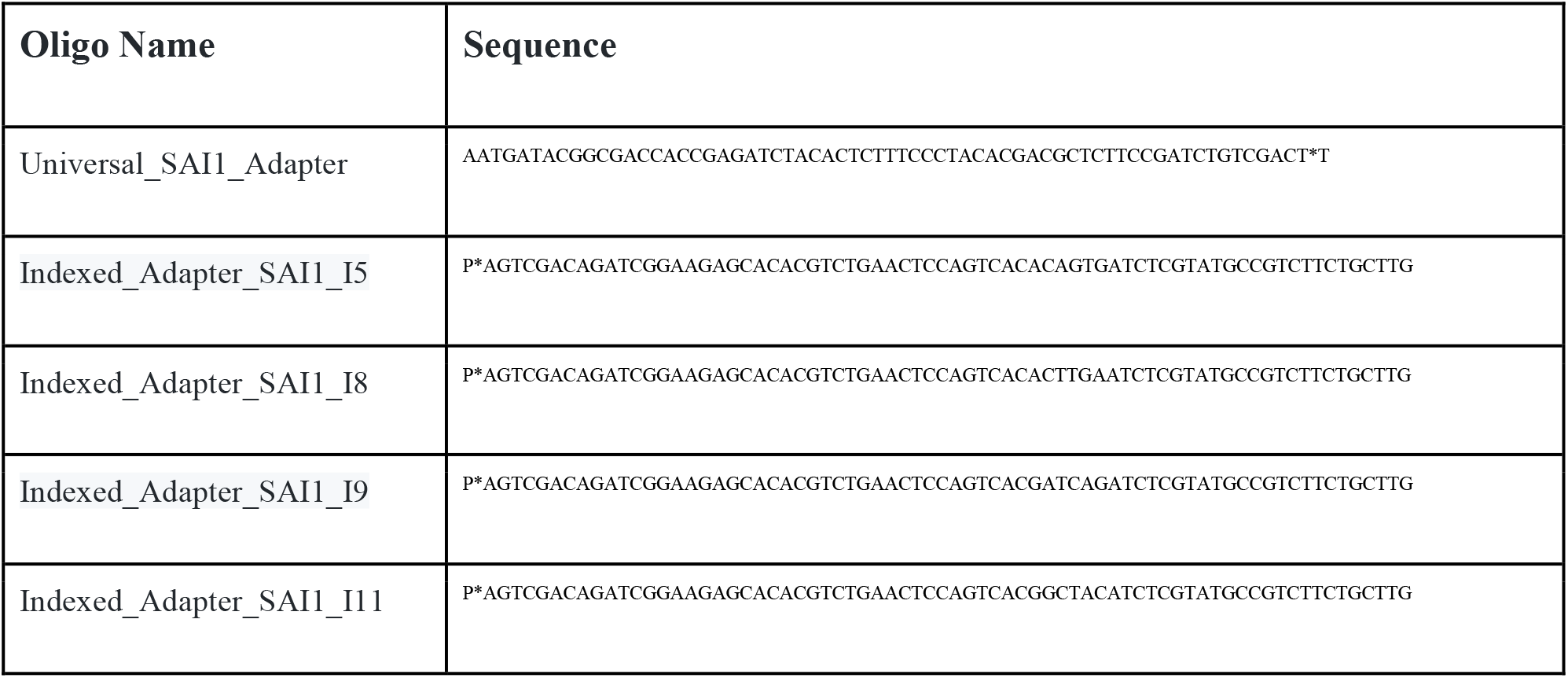
Corrected adapter sequences for mRNA library preparation. Oligos are listed in a 5’ to 3’ orientation with “P” indicates a phosphorylation modification to enable ligation.

*mRNA Library Preparation and Normalization-* Probes were made from two (of four) control individuals and two (of four) exposed individuals. The first step for this subset of individuals was to prepare stranded mRNA libraries using the Kapa Stranded mRNA-Seq Kit (KAPA Biosystems) with the following modifications: custom adapters were used, 4 micrograms of RNA per individual were used as starting material, half volume reactions were used for all steps, adapters were used at a final reaction concentration of 50 nM during ligation, and 12 cycles of PCR were used for enrichment. Complete libraries were visualized on a BioAnalyzer using the DNA 1000 kit, quantified using fluorometry, and then 125 ng of each library was taken and pooled to single library of 500 ng.

To reduce the abundance of highly expressed transcripts in our final probe set, complete libraries were normalized following Illumina’s standard protocol for DSN normalization. First, the cDNA library was heat denatured and slowly allowed to reanneal. Next, the library was treated with duplex-specific nuclease (DSN), which will remove abundant DNA molecules that have properly annealed. After DSN treatment, the library was SPRI purified and enriched via 12 cycles of PCR. A subsample of probes was exposed to an additional 12 cycles of PCR to test for PCR artifacts in probe synthesis. The normalized cDNA library was visualized on a BioAnalyzer using the DNA 1000 kit, quantified with a Qubit DNA Broad Range kit (Invitrogen), and then split into two equal volume tubes, one to be saved for sequencing and one for probe synthesis. The DNS-normalized libraries were sequenced on one half lane of HiSeq 4000 by GENEWIZ (www.genewiz.com).

*Probe Synthesis-To* remove the sequencing adapters, the cDNA library was treated with 100 units of SalI-HF restriction enzyme (New England Biolabs) in a total volume of 40 μl at 37°C for 16 hours. After digestion, the digested library was kept in the same tube, and 4.5 μl of 10X Mung Bean Nuclease Buffer and 5 units of Mung Bean Nuclease (New England Biolabs) were added. The reaction was then incubated at 30°C for 30 minutes. An SPRI cleanup using AMPure XP (Agencourt) was completed with an initial ratio of 1.8X. After, visualization of the library on an Agilent BioAnalyzer, a subsequent SPRI cleanup of 1.5X was completed to remove all digested adapters. The clean, digested cDNA fragments were then biotin labeled using the DecaLabel Biotin DNA labeling kit (Thermo Scientific) using the included protocol. The labeling reaction was then cleaned using a 1.5X SPRI cleanup and fluorometrically quantified. To test the effects of additional PCR cycles on probe effectiveness, 40 ng of the original, normalized cDNA library was subjected to an additional 12 cycles of PCR, and then converted to probes as described above.

*Genomic DNA Library Preparation-* Capture was performed on a standard genomic DNA library. 500 ng of genomic DNA from all eight individuals was sheared to a modal peak of 150 base pairs using a Covaris M220 Focused-ultrasonicator. The sheared DNA was inserted directly into step 2.1 of the KAPA HyperPlus kit with the following modifications: half reaction volumes were used, and a final adapter:insert molar ratio of 50:1 was used with custom TruSeq-style, barcoded adapters (note: the adapters contained erroneous mismatches in the barcodes between the top and bottom oligos; the original oligonucleotide sequences can be found in Supplemental Table 2 and corrected versions in Supplemental Table 3). After adapter ligation, individuals were pooled into one single library, and libraries were enriched with 6 cycles of PCR using primers that complemented the Illumina P5 adapter and Indexed P7 (Supplemental Table 2). The final library was quantified fluorometrically quantified and analyzed on an Agilent BioAnalyzer.

*Hybridization-* Three replicate captures were performed using the set of original probes and the set of probes with 12 extra cycles of PCR. The hybridization protocol closely followed that of Suchan *et al.* (2016). 500 ng of probes and 500 ng of genomic DNA library were hybridized along with blocking oligonucleotides (Table 2) at a final concentration of 20 μM in a solution of 6X SSC, 5 mM EDTA, 0.1% SDS, 2X Denhardt’s solution, and 500 ng o,t-1 DNA. The hybridization mixture was incubated at 95°C for 10 minutes, and then 65°C for 48 hours in a thermocycler. The solution was gently vortexed every few hours.

**Table 2.** Exome capture sequencing, filtering, and mapping statistics. EC_2, EC_4, and EC_7 are the three replicate captures with the original probe pool, and EC_1, EC_3, and EC_12 are the replicate captures with the probe pool exposed to 12 extra rounds of PCR.

| **Replicate**<br>**Capture Pool** | **Raw**<br>**Reads** | **Filtered**<br>**Reads** | **Mapped**<br>**Reads** | **%**<br>**Duplicates** | **Filtered**<br>**Mapped Reads** | **% mapping to**<br>**mitochondrial**<br>**genome** |
| --- | --- | --- | --- | --- | --- | --- |
| EC_2 | 53,493,950 | 53,118,952 | 42,403,525 | 5.8 | 35,955,539 | 1.7% |
| EC_4 | 44,935,340 | 44,651,228 | 35,275,663 | 6.1 | 29,519,347 | 1.6% |
| EC_7 | 43,745,614 | 43,448,296 | 35,007,184 | 5.6 | 29,723,437 | 2.2% |
| EC_1 | 41,402,996 | 41,145,724 | 32,668,750 | 4.5 | 27,940,717 | 1.8% |
| EC_3 | 56,127,536 | 55,753,268 | 44,605,960 | 5.0 | 38,103,268 | 1.9% |
| EC_12 | 46,068,760 | 45,750,394 | 37,227,067 | 6.2 | 31,497,298 | 2.2% |

*Exome Capture-* 40 μl of hybridization mixed was added to 200 μl of DynaBeads M-280 Streptavidin beads (Thermo Fisher Scientific). The beads and hybridization mixture were then incubated for 30 min at room temperature. The mixture was then placed on a magnetic stand until clear, and the supernatant was removed. This was followed by four bead washes under slightly different conditions. First, the beads were washed with 200 μl 1X SSC and 0.1% SSC solution, incubated at 65°C for 15 min, placed on the magnet stand, and the supernatant was removed. Second, the beads were washed with 200 μl 1X SSC and 0.1% SSC solution incubated at 65°C for 10 minutes, placed on the magnet stand, and the supernatant was removed. Third, the beads were washed with 200 μl 0.5 SSX and 0.1% SDS solution, incubated at 65°C for 10 minutes, placed on the magnet stand, and the supernatant was removed. Finally, the beads were washed with 200 μl 0.1X SSC and 0.1% SDS, incubated at 65°C for 10 minutes, placed on the magnet stand, and the supernatant was removed. Lastly, DNA was eluted from the beads in 22 μl of molecular grade water heated to 80°C for 10 minutes. The solution was placed on the magnet and the supernatant was saved. The hybridized fragments were then enriched with 12 cycles of PCR using the appropriate P5 and P7 PCR primers and cleaned with 1X AMPure XP with a final elution in 10 mM Tris-HCl (pH 8.0). The six replicate captures, each containing 8 uniquely barcoded individuals, were sequenced on one half lane (separate from the RNA libraries) on the HiSeq 4000 platform by GENEWIZ (www.genewiz.com).

### Bioinformatic Analysis

All bioinformatic code, including custom scripts and a script to repeat all analyses, can be found at (https://github.com/jpuritz/EecSeq/tree/master/Bioinformatics)

*RNA libraries*-RNA reads were first trimmed for quality and custom adapter sequences were searched for with Trimmomatic (Bolger *et al.* 2014) as implemented in the dDocent pipeline (version 2.2.20; Puritz *et al.* 2014). Reads were then aligned to release 3.0 of the *Crassostrea virginica* genome (Accession: GCA_002022765.4) using the program STAR (Dobin *et al.* 2013). The genome index was created using NCBI gene annotations for splice junctions. Reads were aligned in a two-step process, first using the splice junctions in the genome index, and then again using both the splice junctions in the index and additional splice junctions found during the first alignment. Alignment files from the four libraries were then merged with SAMtools (version 1.4; Li *et al.* 2009) and filtered for MAPQ > 4, only primary alignments, and reads that were hard/soft clipped at less than 75 bp. SAMtools (Li *et al.* 2009) and Bedtools (Quinlan 2014) were used to calculate read and per bp coverage levels for exons, introns, and intergenic regions.

*EecSeq Libraries*-Raw reads were first trimmed using the standard methods in the dDocent pipeline (version 2.2.20; Puritz *et al.* 2014). The DNA adapters contained erroneous mismatches between the top and bottom oligos in the barcode (original oligonucleotide sequences can be found in Supplemental Table 2 and corrected versions in Supplemental Table 3). These differences prevented demultiplexing beyond the capture pool level, and also lead to potentially erroneous base calls within the first 7 bp of sequencing. To remove these artifacts, the first 7 bp of every forward read were clipped. Additionally, adapter sequences were searched for in the paired-end sequences using custom scripts. After trimming, reads were aligned to the reference genome using BWA (Li & Durbin 2009) with the mismatch parameter lowered from 4 to 3, and the gap opening penalty lowered from 6 to 5. PCR duplicates were marked using the *MarkDuplicatesWithMateCigar* module of Picard (http://broadinstitute.github.io/picard), and then SAMtools (Li *et al.* 2009) was used to remove duplicates, secondary alignments, mappings with a quality score less than ten, and reads with more than 80 bp clipped. SAMtools (Li *et al.* 2009) and Bedtools (Quinlan 2014) were used to calculate read and per bp coverage levels for exons, introns, and intergenic regions. FreeBayes (Garrison and Marth 2012) was used to call SNPs.

*Calculating Capture Efficiency-* EecSeq is unique amongst exome capture methods because the probes are not designed directly, i.e. there is no set of *a priori* targets. Additionally, EecSeq is designed to capture exons that are expressed in the samples used to create probes - not the entire exome. To compare EecSeq to other capture methods, capture targets were defined as exons that had more than 35X coverage in the RNAseq (probe) data and confidence intervals were generated by defining capture targets as 20X RNAseq coverage and 50X RNAseq coverage. We also calculated a conservative, near-target range of 150 bp on either side of the defined targets. This range corresponds to the modal DNA fragment length used for the capture libraries with the expectation that exon probes could capture reads that far from the original target.

## Results

*RNA sequencing results*-RNA sequencing, filtering, and mapping statistics can be found in Supplemental Table 3. After filtering, a total of 21,990,025 RNA reads were mapped uniquely to the eastern oyster genome. Of the total RNA reads, 78% mapped to genic regions of the genome, and 58% mapped to annotated exon regions. Across all exonic bases in the genome, less than 5% had more than 50X coverage; however, over 16% had at least 20X coverage and over 45% had at least 5X coverage (Figure 2).

**Figure 2.**
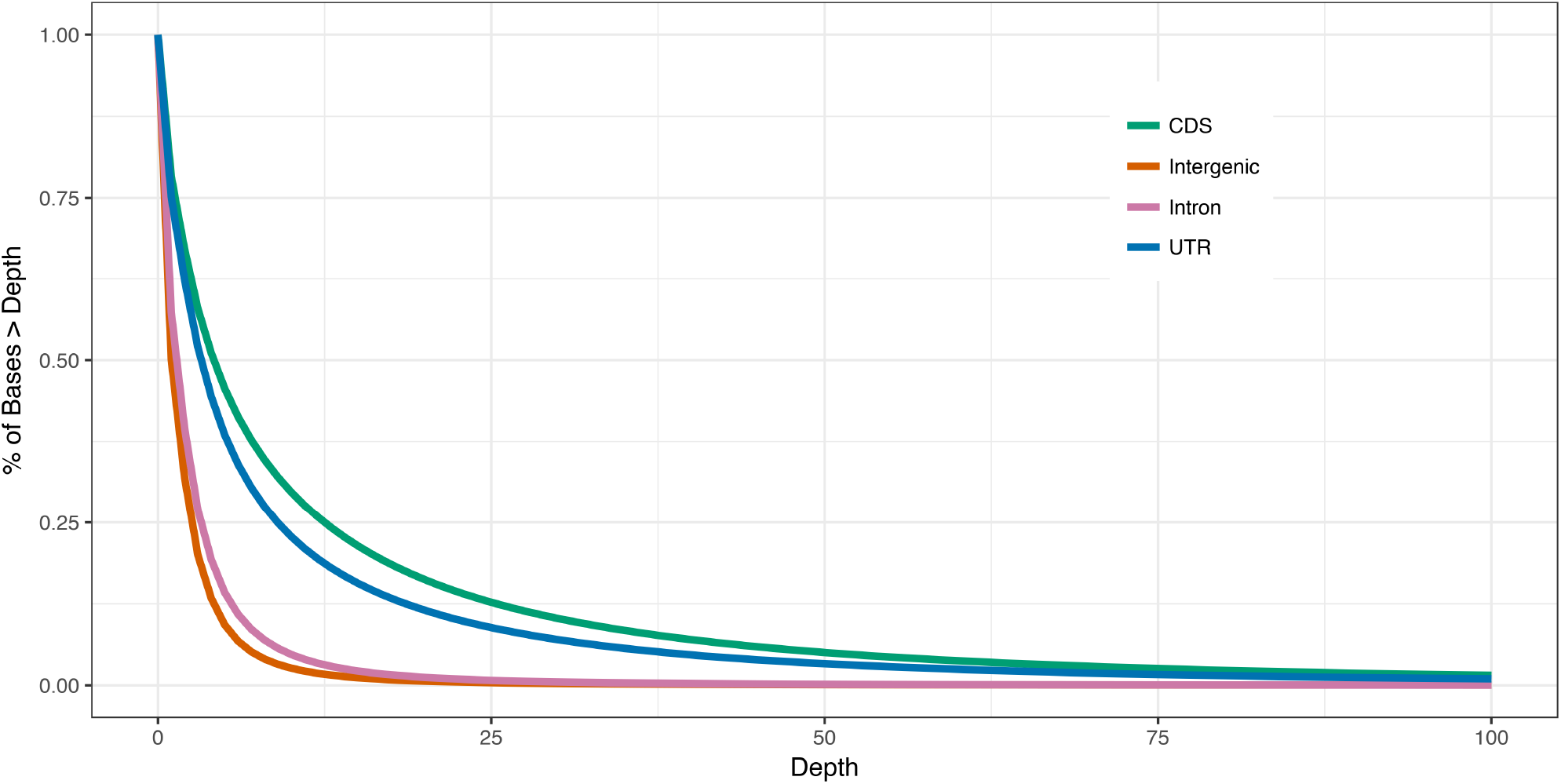
Distribution of RNA reads across regions of the oyster genome. Percentage of bases within exons-both coding sequences (CDS) and untranslated exon regions (UTR), intergenic, and intron regions at various coverage levels.

*Exome capture sequencing results-* Six replicate capture pools of the same eight individuals were sequenced on half a lane of Illumina HiSeq (3 replicates from probes that had been enriched via 12 cycles of PCR and 3 replicates from probes that had been enriched via 24 cycles of PCR). A summary of exome capture sequencing, filtering, and mapping statistics are shown in Table 2. On average, there were 47,629,033 raw reads (forward and paired-end) per capture pool and an average of 32,123,268 mapped reads per capture pool after filtration. Across the entire oyster genome, RNA sequencing coverage and exome sequencing coverage was highly correlated (Supplemental Figure 1), and across all exon regions total RNA coverage predicted 72.6% of the variation in exome capture coverage (Figure 3; log-log transformation, *R^2^* = 0.72619, *p* < 0.0001). Coverage across all exons and expressed exon targets was highly correlated (0.984 < *r* < 0.996) across all replicate captures, and the average capture of pools with standard probes and the average capture of pools with probes with extra PCR was virtually identical (R^2^ = 99.1; *p* < 0.0001).

**Figure 3.**
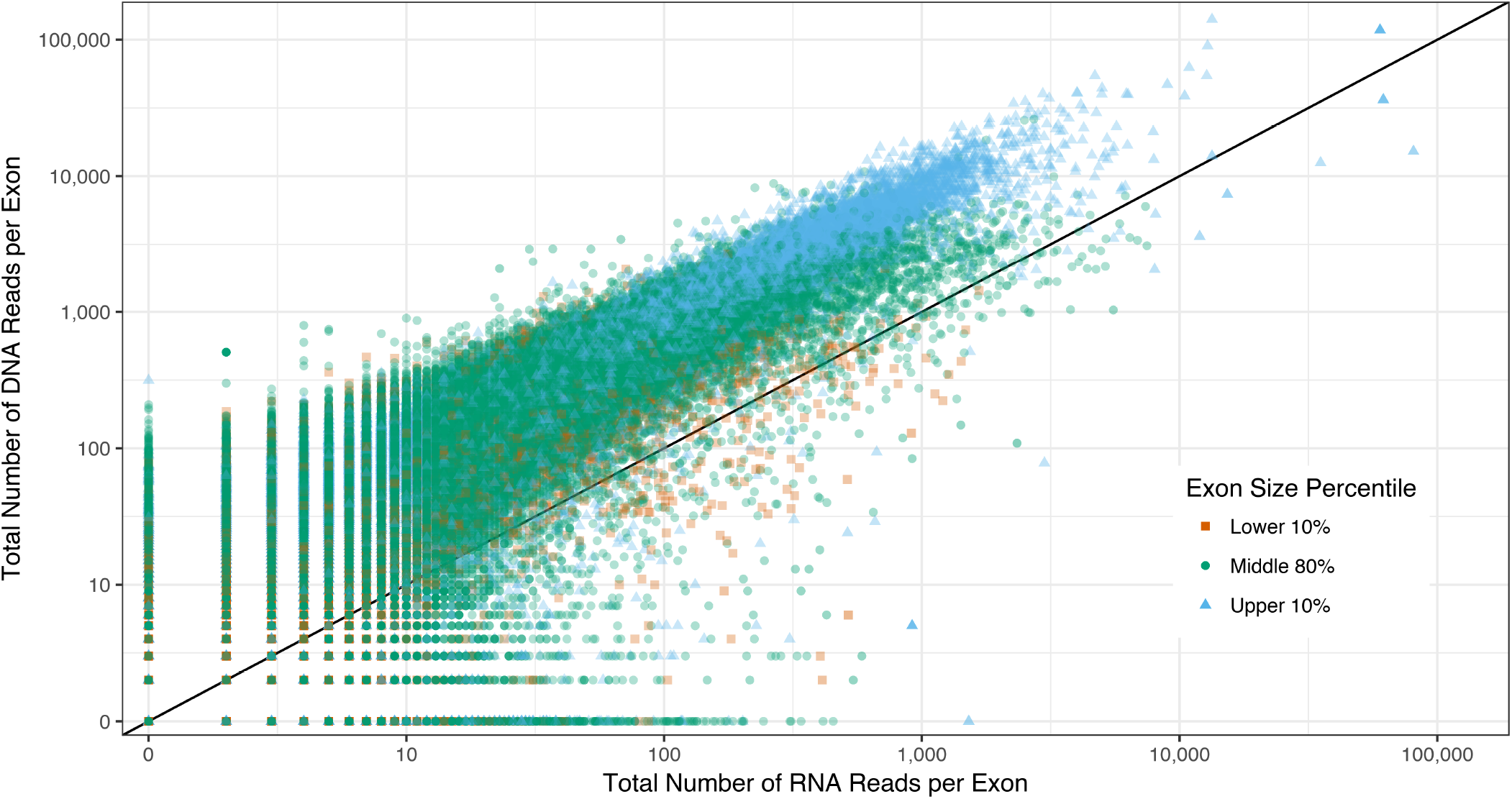
Total DNA and RNA coverage across all exons. Depth was calculated as the total number of reads overlapping with an exon region. For exome capture depth (y-axis), reads were summed across all 6 replicate captures. For RNA read depth, reads were summed across all four libraries. The shape and color of each point was determined by the percentile size of the respective exon (lower 10% < 59 bp, upper 10% > 517 bp, and the middle 80% was between 57 bp and 517bp). Note that the DNA reads were sequenced to greater depth than the RNA-derived probes.

*Exome capture efficiency-* Capture sensitivity, or the percentage of targets covered by at least one read (1X), was high across all replicate pools, regardless of target set (Table 3). Across all known exons, sensitivity was on average 86.8% across replicate capture pools, and across all defined target sets, sensitivity was over 99.4%. Increasing the sensitivity threshold from 1X to 10X lowers the sensitivity across all exons but has little effect on sensitivity across defined target sets (Supplemental Table 4). Sensitivity can also be measured at the per bp level instead of per exon. The percent of target bases captured is shown as a function of sensitivity threshold (read depth of capture libraries) in Figure 4.

**Table 3.**
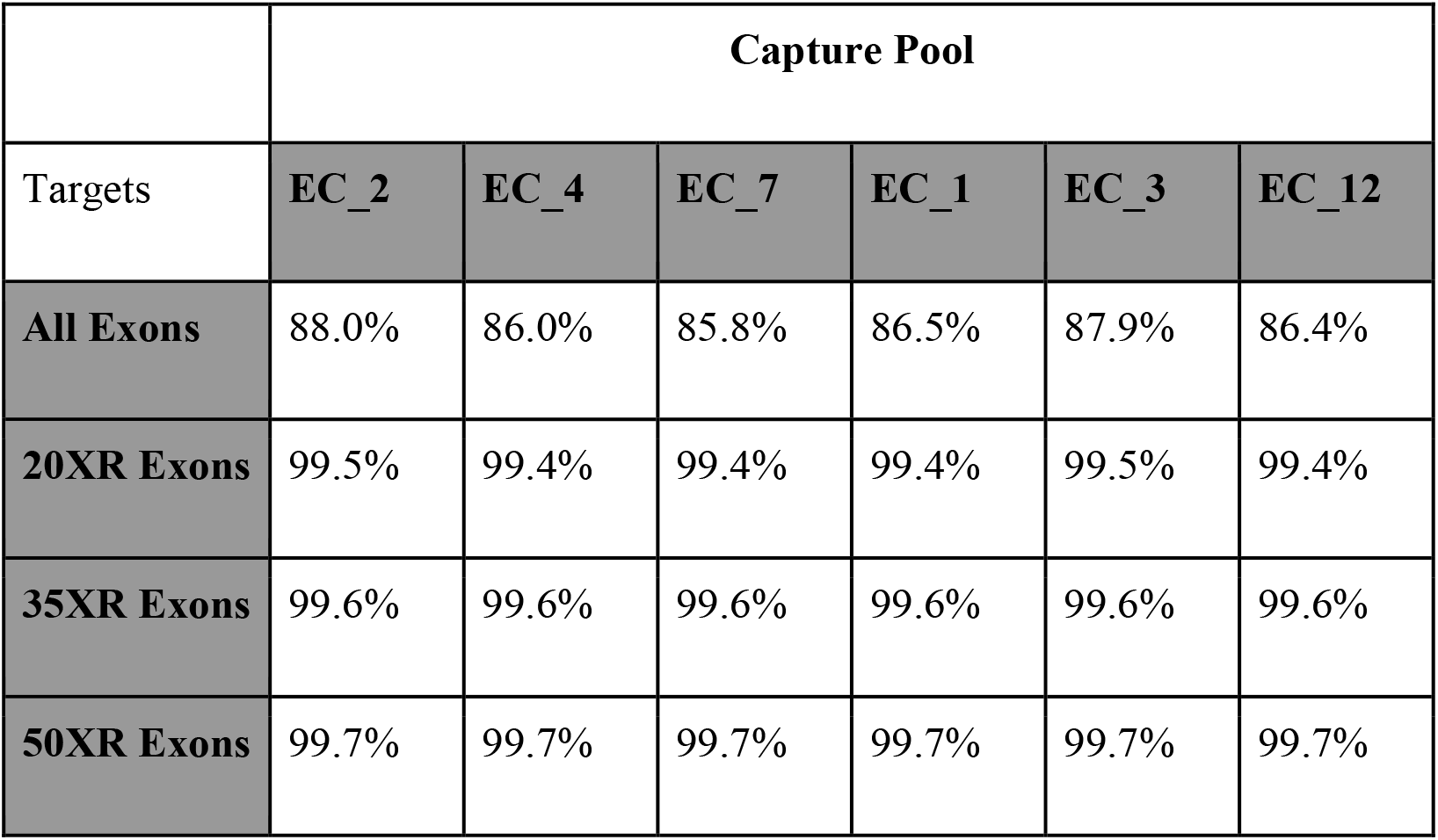
Exome capture sensitivity with a 1x threshold. Sensitivity is the percentage of target bp with at least one read mapping successfully. Here, targets are broken up into subsets: All annotated exons, exons with at least 20X coverage from the RNA library, exons with at least 35X coverage from the RNA library, and exons with at least 50X coverage from the RNA library. EC_2, EC_4, and EC_7 are the three replicate captures with the original probe pool, and EC_1, EC_3, and EC_12 are the replicate captures with the probe pool exposed to 12 extra rounds of PCR.

**Figure 4.**
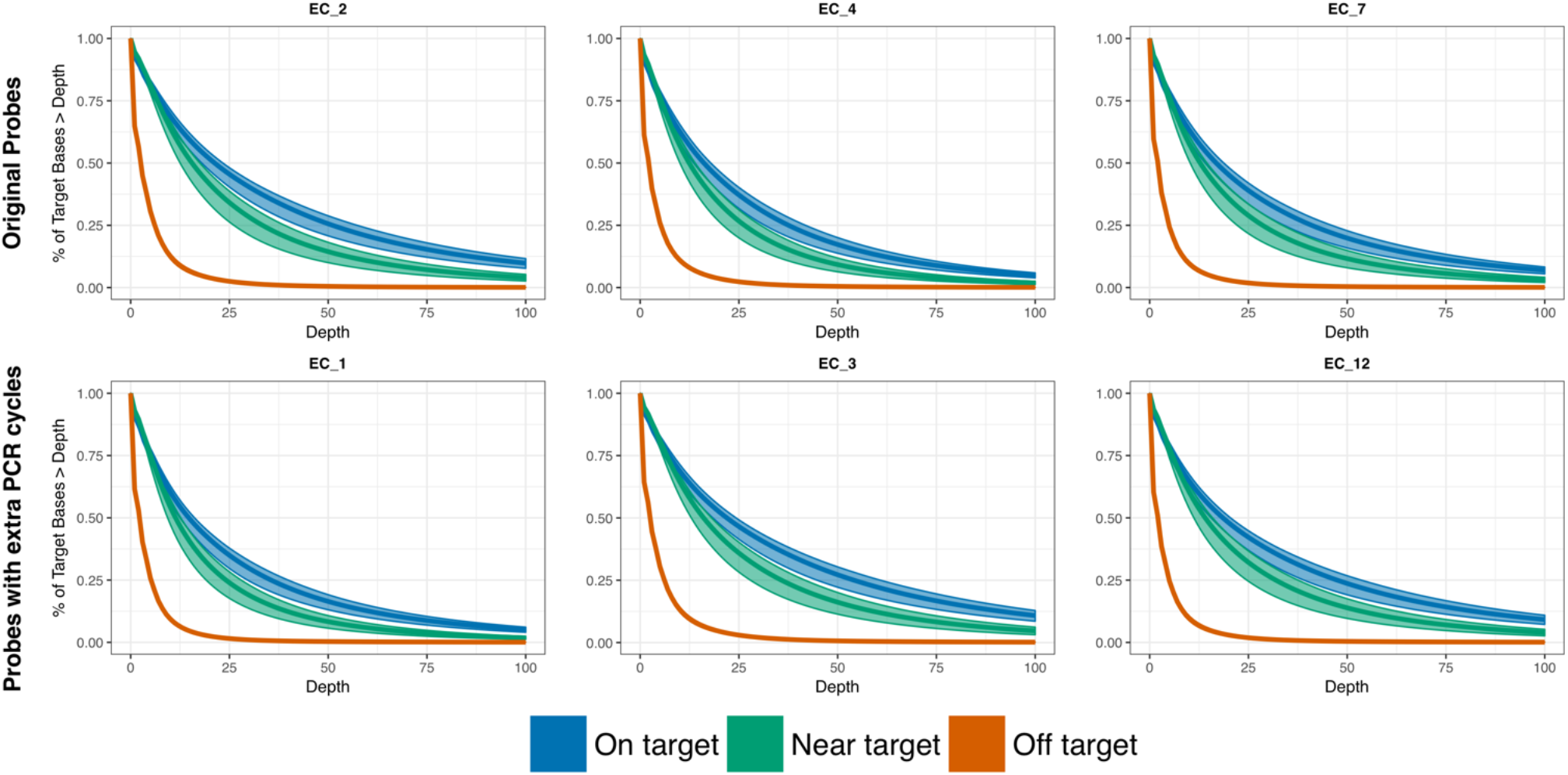
Per base pair EecSeq capture sensitivity. To measure EecSeq capture (DNA) sensitivity, capture targets were defined as exons that had more than 35X coverage in the RNAseq (probe) data. Confidence intervals were generated by defining capture targets between 20X RNAseq coverage and 50X RNAseq coverage. Near-target mapping were 150 bp on either side of the defined targets. This range corresponds to the modal DNA fragment length used for the capture libraries with the expectation that exon probes could capture reads that far from the original target. EC_2, EC_4, and EC_7 are the three replicate captures with the original probe pool, and EC_1, EC_3, and EC_12 are the replicate captures with the probe pool exposed to 12 extra rounds of PCR. Depth in this figure is the depth of DNA reads from EecSeq captures.

Capture specificity is the percentage of mapped reads that fall within target regions. Across all exons, capture pools averaged 47.9% reads on target, 6.8% of reads near target (falling within 150 bp of an exon, one modal read length), and 45.3% of reads off-target (more than 150 bp away from an exon). Across defined expressed exon targets (exons that sequenced to 35x read depth), capture pools averaged 37.1% (C.I. 33.6% - 41.4%) reads on target, 3.55% (C.I. 3.0% - 4.4%) of reads near target, and 59.38% (C.I. 54.2% - 63.4%) reads off target.

For all exons, between the 10th and 90th percentile of exon length (59bp - 517bp), the mean per basepair coverage averaged 17.75X +/- 0.06X for each capture pool of 8 individuals. When considering target exons (35X coverage in RNA-derived probes), the mean per basepair coverage increased to 61.22X +/- 0.23X on average for each capture pool. This breaks down to approximately 7.66 reads on average per individual per bp within expressed exome targets. Within exons, mean per basepair coverage was evenly distributed across all base pairs with only slightly lower coverage at the 5’ or 3’ edges of exons compared to the middle of exons (Figure 5; Supplemental Figure 3).

**Figure 5.**
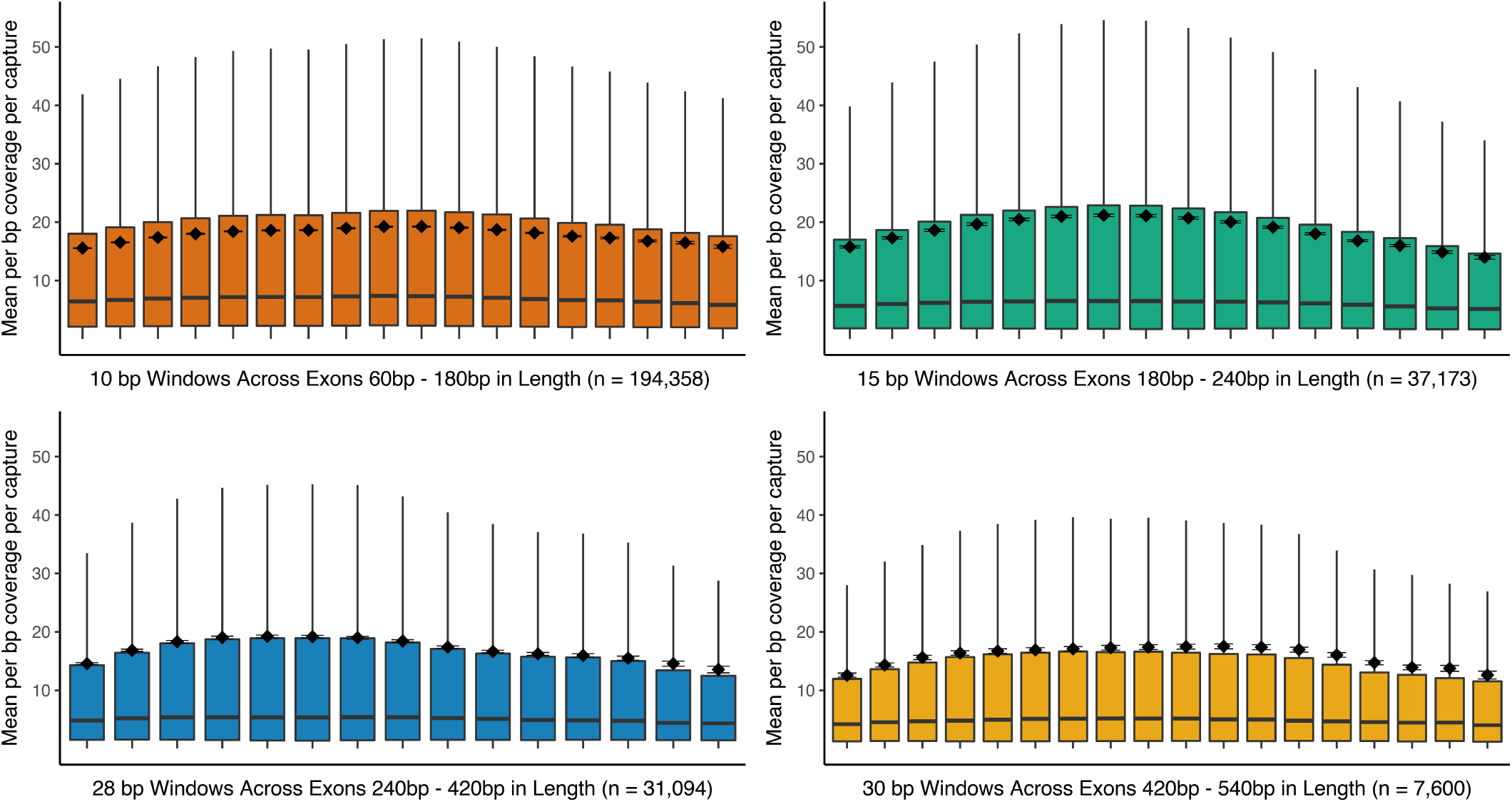
Boxplots of mean per basepair coverage levels plotted across exons size windows. All annotated exons were broken into 10bp - 30 bp windows depending on overall size and the mean per basepair coverage per capture was calculated for each window size. The line each box represents the median of mean coverage values and the box surrounds the 25^th^ and 75^th^ percentiles. The mean of each bin class is plotted as a black diamond with standard error bars around it. Outlier points were not plotted. Note that the data for this graph is for all annotated exons, regardless of expected capture. See Supplemental Figure 3 for a similar plot focused on an expressed target set.

Mean capture coverage also did not appear to relate to the GC content of the target exon (Figure 6), though it did appear to peak near the mean GC content of 43.57%. To test this, we calculated the reciprocal of the absolute value of the difference between each exon GC content and the average GC content, and then tested for a linear relationship to mean coverage. Though we found this relationship to be significant (*p* > 0.0008), it explained only the 0.0033% of the variance in coverage, confirming that exon GC content did not affect exon capture in a meaningful way.

**Figure 6.**
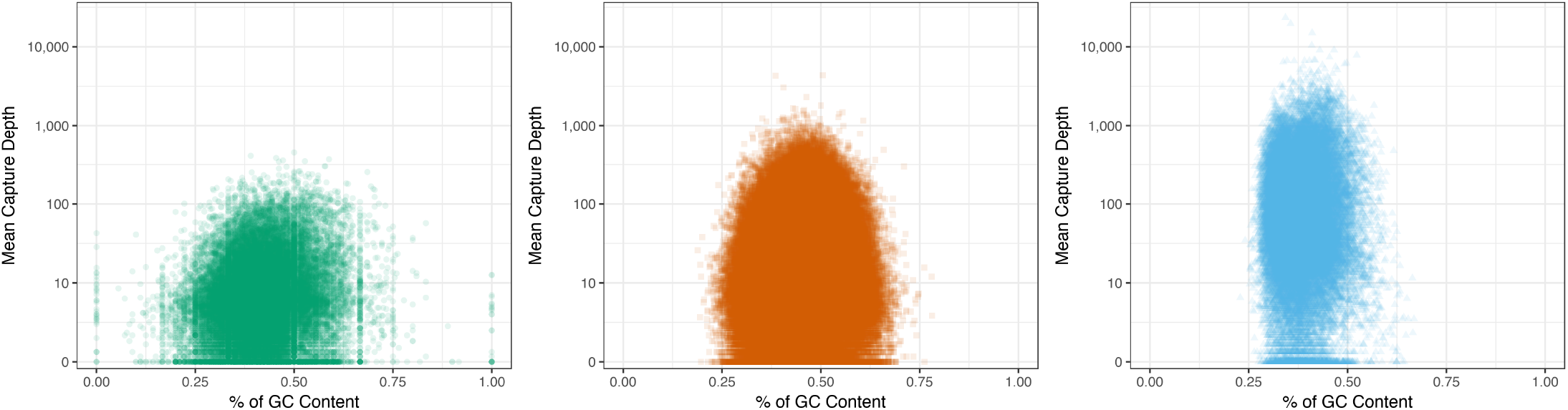
Mean capture depth plotted against exon GC content. Exons were broken up into three size windows: (1) Lower 10%- exons less than 57 bp, (2) Middle 80%- exons greater than 56 bp and less than 518, (3) Upper 10%- exons greater than 517 bp.

Coverage did vary significantly between untranslated regions (UTR) within exons and coding sequence (CDS) within exons (Welch’s test *t* = 40.063; degrees of freedom = 135580; *p* < 0.0001) with a mean coverage for UTR equaling 11.59X +/− 0.0864 and a mean coverage for CDS equaling 17.71X +/− 0.1261. This small but significant coverage difference was also evident as the percent of target bases greater than a given read depth (Supplemental Figure 2). This pattern was not surprising, however, because the same pattern was observed for the RNA reads (CDS mean coverage = 13.65X +/− 0.2011; UTR mean coverage = 8.25 +/− 0.1275; Welch’s test *t* = 22.677; degrees of freedom = 129300*;p* < 0.0001), indicating that the probes also had lower coverage in UTR compared to CDS.

*Expressed exon capture*- To visualize the relationship between coverage and an expected expressed target, we plotted coverage of the six capture pools along two heat shock proteins, Heat Shock cognate 71 kDa (NCBI Reference Sequence: XM_022472393.1, Figure 7) and Heat Shock 70 kDa protein 12B-like (NCBI Reference Sequence: XM_022468697.1; Supplemental Figure 4). As expected, exons in both genes show elevated coverage that corresponded to the coverage of the mRNA-derived probes, especially along regions with corresponding CDS with few reads mapping to intronic or intergenic regions.

**Figure 7.**
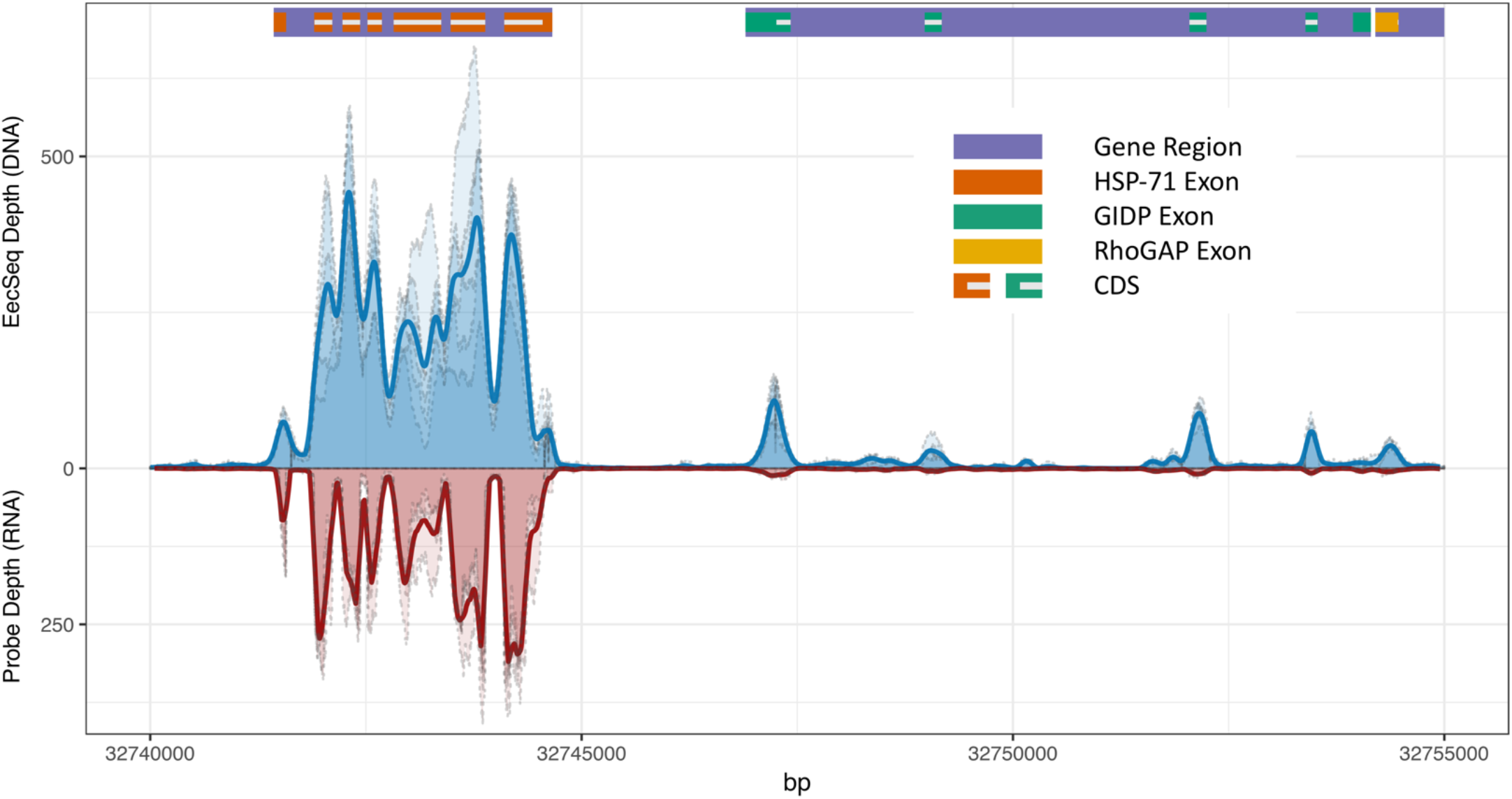
EecSeq capture and probe coverage across Heat Shock cognate 71 kDa. Coverage for each replicate capture pools is plotted along base pairs 32,740,000 to 32,755,000 of reference Chromosome NC_035780.1 containing the full gene region of Heat Shock cognate 71 kDa (NCBI Reference Sequence: XM_022472393.1), predicted glucose-induced degradation protein 8 homolog (NCBI Reference Sequence: XM_022486802), and a partial gene region for rho GTPase-activating protein 39-like (NCBI Reference Sequence: XM_022486743.1). Each exome capture pool coverage is plotted in light blue with dashed grey border, and a rolling 100 bp window average across all pools is plotted in dark blue. Each RNAseq (probe) sample coverage is plotted in light red with dashed grey border and a rolling 100 bp window average across all pools is plotted in dark red. Gene regions are marked in purple with exons color coded by gene. Coding sequence (CDS) is marked by a white bar within exon markers.

*SNP Discovery-* A total of 1,011,107 raw SNPs were discovered with 909,792 SNPs having a quality score higher than 20. A total of 99,169 high quality SNPs were found within known exons. Of these, 31,579 exome SNPs had at least an average of 16X coverage, 15,760 exome SNPs had at least an average of 32X coverage, 8,837 exome SNPs had at least an average of 48X coverage, and 3,508 exome SNPs had at least an average of 80X coverage with an additional 2,443 80X-SNPs found outside of exon regions.

## Discussion

Expressed exome capture sequencing (EecSeq) is a novel design for exome capture that uses *in-situ* synthesized biotinylated cDNA probes to enrich for exon sequences, thereby removing the requirement of *a priori* genomic resources, costly exon probe design, and synthesis. Here, we showed that EecSeq target enrichment had high levels of sensitivity, with comparable if not superior performance and specificity to traditional methods. EecSeq exon enrichment showed even coverage levels with exons and across exons with differing levels of GC content. Lastly, we showed that EecSeq can quickly and cheaply generate thousands of exon SNPs.

### Benefits of EecSeq

*Diverse probes-* With EecSeq, cDNA exon probes are constructed *in-situ* from extracted mRNA, and this allows for the design of a high-diversity probe pool. Traditional sequence capture probes are typically designed from a single reference genome or individual, and this may limit capture efficiency on individuals with different SNPs, insertions, or deletions than the reference. While probes been successfully used to capture sequences in quite divergent species (less than 5% sequence divergence, Jones & Good 2016), there is evidence that capture success declines as sequences become less related to the reference. Portik *et al.* (2016) found that for each percent increase of pairwise divergence, missing data increased 4.76%, sensitivity decreased 4.57%, and specificity decreased 3.26%. Even with well-designed, commercially available capture kits for human exon capture, Sulonen *et al.* (2011) found that allele balances for heterozygous variants tended to have more reference bases than variant bases in the heterozygous variant position across all methods for probe development. Insertions and deletions (InDels) are arguably an even larger problem, since these would decrease hybridization with a probe due to a frameshift.

*Longer Probes-* Traditional exome capture relies on synthesized RNA or DNA baits. These baits can be relatively small (60 bp; Bi *et al.* 2012) or range between 95 and 120 bp (Clark *et al.* 2011; Sulonen *et al.* 2011; Nadeau *et al.* 2012; Chilamakuri *et al.* 2014). In contrast, EecSeq probes have a modal length of 150 bp but also range up to over 400 bp (data not shown). The longer length of EecSeq probes likely helps to buffer against divergence between probes and targets. The longer probes may also be the reason why we observed relatively little GC bias in coverage across exons, and may help explain the uniformity of coverage within exons in EecSeq data.

*Cost-* EecSeq provides significant cost and time savings over traditional exome capture and RNA sequencing (RNAseq). No *a priori* genomic information is necessary for EecSeq, saving substantial time and money for obtaining these data in non-model organisms. Likewise, the cost of synthesizing the probes is significantly reduced because probes can be made in-house and do not have to be designed by a company. On a per sample basis, EecSeq is also significantly cheaper than RNAseq because (i) commercial DNA library preps are cheaper than those for mRNA, and (ii) more individuals can be multiplexed on a single lane. For example, the cost of RNA seq is $246 per sample (cost estimated using the same RNA kits used with EecSeq and ½ reactions) and assuming that 12 RNAseq libraries can be sequenced in a single lane of Illumina HiSeq, the cost per sample is ($1,008; cost of the kit; Kapa Biosystems Stranded mRNA-Seq Kit with 24 reactions or 48 half-reactions)*(1/48; the amount used per sample) + $2700/12 = $246 per sample). The equivalent cost per sample for EecSeq is $48.02 per sample (for 96 samples in one lane of HiSeq; Supplemental Table 6) or $62.08 per sample if a more conservative sequencing strategy is used (96 samples sequenced over 1.5 lanes of HiSeq; Supplemental Table 6).

*No dependency on restriction sites*- A recently published method, hyRAD-X, (Schmid *et al.* 2017) is similar to EecSeq in that it uses *in-situ* synthesized cDNA probes from expressed mRNA to capture exome sequences. However, the protocol relies on a restriction digest to fragment cDNA and ligate on probes. This may result in a reduced template of probes because not all cDNA fragments will have restriction sites on both ends. To evaluate the possibility that the hyRAD-X would produce a reduced template of probes, we performed crude calculations using SimRAD in R (Lepais & Weir 2014) on the *C. virginica* exome. Of the 31,383 known mRNA transcripts in the oyster genome (assuming 1 transcript variant), 29,555 contain at least 2 MseI cut sites (TTAA). However, there is an SPRI cleanup on the digestion (2X), meaning that at best, only fragments 100bp and larger are getting through to biotinylation

(http://www.keatslab.org/blog/pcrpurificationampureandsimple). SimRAD estimates 220,184 out of a possible 440,881 fragments. Therefore, at the absolute best hyRAD-X is only sampling (29,555/31,383)*(220,184/440,881) = 47% of the exome, though this number may increase slightly due to transcript variations. Relying on restriction digests may also produce skewed size distributions in probes which would be magnified in subsequent rounds of PCR. In Schmid *et al.* (2017), hyRAD-X generated 524 exome SNPs at a minimum of 6X coverage across 27 samples (compared to the 3,508 exome SNPs discovered at 80X coverage derived from only 8 effective samples in 6 replicate capture using EecSeq), but they were also studying ancient DNA and so whether the hyRAD-X protocol results in limited coverage across exons remains to be tested.

### Caveats of EecSeq

Despite the demonstrated benefits of EecSeq, there are some potential caveats that should be considered before employing the method. First, there is no ability to filter out probes that belong to repetitive sequences, which are often present at high concentrations in large-genome organisms such as amphibians (Keinath *et al.* 2015) or conifers (De La Torre et al. 2014). In one capture study from designed probes, a small proportion of the probes (unknowingly at the time of probe development) matched highly repetitive sequences (Syring *et al.* 2016). This resulted in an inordinate number of reads to these few probe sequences (Syring *et al.* 2016). However, the inclusion of known repetitive sequence blocker in hybridization, such as c_0_t-1 that is used in the EecSeq protocol, has been shown to nearly double capture efficiency (McCartney-Melstad *et al.* 2016). In general, repetitive elements, short repeats, and low complexity regions are problematic for all types of probe design and capture.

Another caveat of using EecSeq is the need to obtain RNA from relevant samples, although capture designs or gene expression studies based on transcriptomes face the same challenge. Note, however, the advantage that EecSeq probes can be made from mRNA pooled from many individuals, tissues, and conditions of interest. If genes of interest are expressed in tissues that are difficult to dissect or are in small abundances (such as neurons), then the RNA-based methods presented here would not be a feasible approach unless pooling multiple extractions.

Additionally, the probes are a limited resource - our results indicate, however, that additional rounds of PCR on the probes have little effect on capture.

### Unique Aspects of EecSeq

Our approach relies on expressed mRNA for probe synthesis and the abundance of particular mRNAs will vary depending on gene expression. EecSeq includes a normalization step to decrease the abundance of very common transcripts, but probe pools will still skew towards highly expressed genes and therefore capture coverage will be higher for those exons. This aspect of EecSeq can be customized for particular research questions. For projects focused on total exome capture, pools from multiple individuals, tissue types, and environmental/laboratory exposures can be constructed to generate a robust probe set. On the contrary, if an investigator is focused on a subset of genes that are responding to a particular stressor, it is possible to make probes from organisms exposed that specific condition and then use those probes to capture other individuals. This reduced probe set may also allow for greater multiplexing, but this remains to be specifically tested. While we have only used mRNA to create probes, there may be possibilities to capture other types transcribed sequences such as long non-coding RNAs or possibly even miRNA.

Previous work on exome capture probe design has focused on intron/exon boundaries. In general, it is thought that capture probes that span exon boundaries will result in low coverage of these regions (Jones & Good 2016) or that certain regions will not be covered at all (Neves *et al.* 2013). Inclusion of too many boundaries may also lower overall capture performance by increasing off-target capture (Suren *et al.* 2016). EecSeq exome probes are derived from mature RNA, so some of the probes will span inevitably span exon boundaries. Though exon/intron boundaries cannot be eliminated in EecSeq, both input mRNA and genomic DNA were fragmented down to a modal size of 150 base pairs, with the intention of making both smaller than the average exon size of Eastern Oysters (note that this size is at the lower limit of what is possible with Illumina sequencing). We found that coverage within exons was fairly uniform, indicating a lack of “edge effects.” We hypothesize that the relative long length of EecSeq probes (compared to commercially synthesized probes), the near matching length of genomic DNA fragments, and the length distribution relative to actual exon size helped to ensure uniform exon coverage.

We compared our observed measures of sensitivity and specificity to other recently published studies in non-model species where probes were designed from bioinformatic resources for the same species. EecSeq capture efficiency performed as well as or outperformed almost all other previously published exome capture studies in non-model species (excluding mice and humans; Table 4) with the notable exception of black cottonwood (Zhou & Holliday 2012), a species with exceptional genomic resources. Note, however, that we analyzed capture efficiency across pools of 8 individuals, and there could be considerable variability at the individual level that remains to be quantified.

**Table 4.** Comparing specificity and sensitivity across capture methods. A summary of sensitivity and specificity of recent exome-capture studies in which probes were designed from the same species. NR: not reported.

| Reference and species | Num. target genes or exons | Sensitivity % of targeted regions > 10x depth | Specificity % of reads mapping to targeted bases | % of reads mapping near target | % of reads mapping off target | Notes |
| --- | --- | --- | --- | --- | --- | --- |
| EecSeq (this study)<br>eastern oyster<br>*Crassostrea virginica* | 71,105<br>(51,096-110,020) | All exons: 54.7%<br>Expressed Exons: 98.8% ( 97.4% - 99.1%) | All exons: 47.8%<br>Expressed Exons: 37.0% ( 33.6% - 41.4%) | All exons: 28.4%<br>Expressed Exons: 23.6% (22.3% - 25.2%) | All exons: 23.7%<br>Expressed Exons: 39.3% (33.3% - 44.1%) |  |
| (Suren *et al.* 2016)<br>pine and spruce<br>*Picea glauca x engelmannii*<br>and<br>*Pinus contorta* | 26824 genes<br>(pine) 28649 genes(spruce) | 51% (spruce) and 59% (pine) (all samples, also metrics for 75% of samples) | 18.5% (spruce) and 21 % (pine) | 37% (spruce) 38% (pine) | 44% (spruce) and 41% (pine) | Non-model species, large genomes, near target defined as 500 bp |
| (Zhou & Holliday 2012)<br>black cottonwood<br>*Populus trichocarpa* | 20.76Mb (5%) of exons, regulatory regions | 86.8 % (at 100X coverage about 0-8%) | ~93% | On average, approximately 80 base pairs nearest the bait were sequenced at a depth of > 10X | NR | Model species with good genome. Off target defined as > 250bp away. |
| (Hebert *et al.* 2013)<br>lake whitefish<br>*Coregonus clupeaformis* | 11,975 nuclear exons, and other genomic markers using 62,438 probes | NR | 11.8% | NR | NR | 98% of targeted genes (2728) were successfully captured a mean read depth of 31X |
| (Bi *et al.* 2012)<br>chipmunk<br>*Tamias alpinus* | 11,975 exons | 40.3% | 25% | NR | NR | % of exons that were covered by at least one read, > 99% |
| (Christmas *et al.* 2017)<br>narrow-leaf hopbush<br>*Dodonaea viscosa* ssp. *angustissima* | 700 genes | NR | 15.7% | NR | NR | Did not account for intron sites |
| (Syring *et al.* 2016)<br>whitebark pine *Pinus albicaulis* | 7,849 distinct transcripts | NR | 13% | NR | NR |  |
| (Müller *et al.* 2014)<br>douglas-fir<br>*Pseudotsuga menziesii* | 57,110 exons | 90% | 32-52% per individual | NR | NR |  |
| (Nadeau *et al.* 2012)<br>butterflies | BAC loci (3.5 MB; 57,610 baits) | 75.6% | 33.5% | NR | NR |  |

### Conclusions and Future Directions

Here, we have shown that EecSeq effectively targets expressed exons, delivers consistent and efficient exome enrichment that is comparable to traditional methods of exome capture, and generates thousands of exome-derived SNPS cost effectively. Additional tests are needed to examine the efficiency of exome capture across individuals for different species, which should be coupled with sequencing of EecSeq probes to investigate the effects of probe pool diversity and sequence divergence between probes and targets on capture. Nonetheless, EecSeq holds substantial promise as a universally applicable and cost-effective method of exome sequencing for virtually any macroscopic organism.

## Acknowledgements

The authors would like to thank Alan Downey-Wall and Sara Schaal for informative discussions and input throughout the duration of this project. The authors also thank Nadir Alvarez for thoughtful comments on a preprint of this manuscript. This work was funded with funds provided to KEL from Northeastern University and NSF OCE-1635423.

## Data Accessibility

Raw, demutliplexed sequences are archived at the NCBI Short Read Archive (BioProject: PRJNA423022). A complete and updated EecSeq protocol can be found at (https://github.com/jpuritz/EecSeq) along with bioinformatic code to repeat all analyses described in this paper.

## Author Contributions

JP conceived the original concept of this work and performed all laboratory and data analysis.

KL contributed all reagents and experimental materials. JP and KL designed the research, experiments, and data analysis, and wrote the manuscript.

## Tables

**Supplemental Table1:**
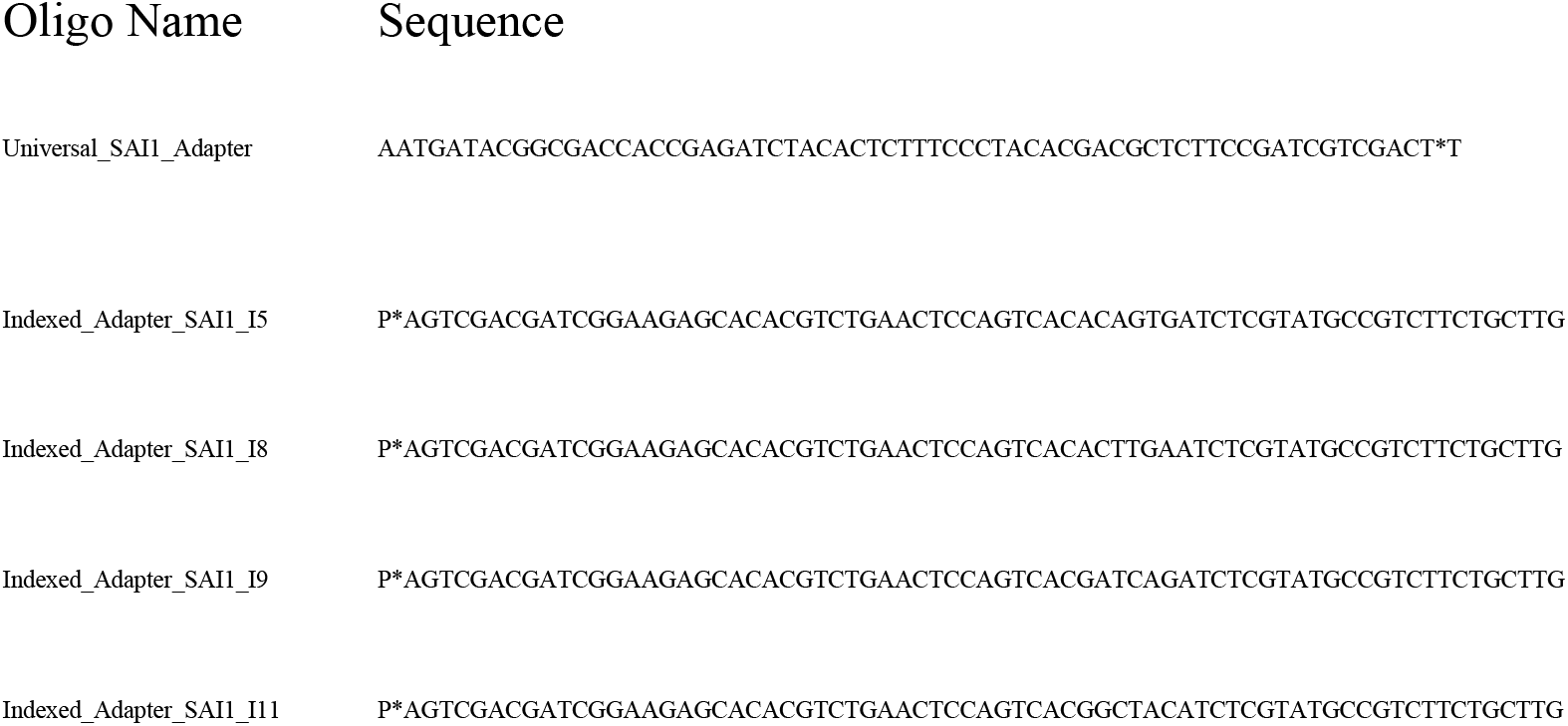
Original RNA Adapters.

**Supplemental Table2:**
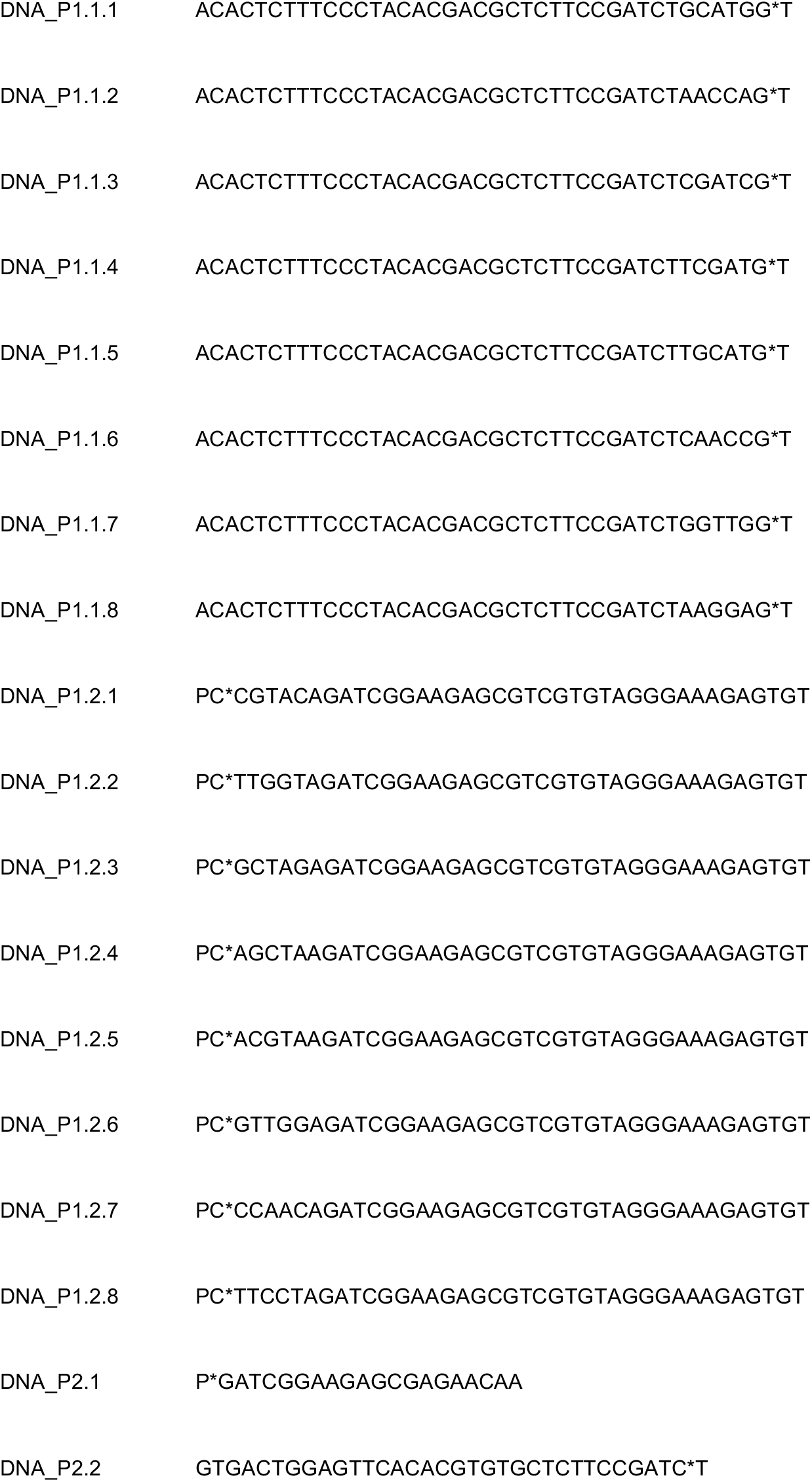
Original DNA Adapters. Oligos should be paired for adapter formation following 1.1. X pairs with 1.2.X.

**Supplemental Table3:**
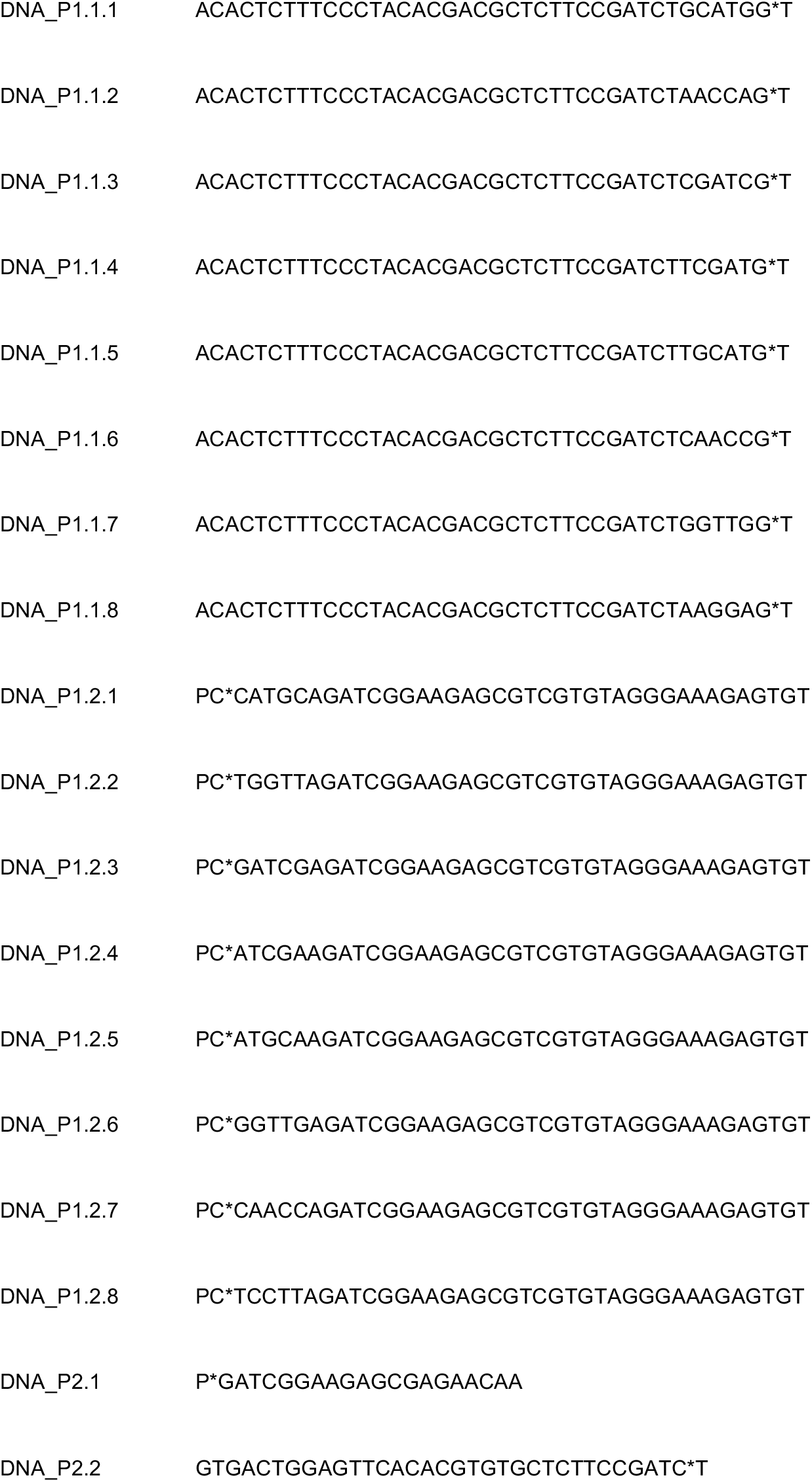
Correct DNA Adapters. Oligos should be paired for adapter formation following 1.1. X pairs with 1.2.X.

**Supplemental Table 4.** RNA sequencing statistics. Samples 3E and 4E were heat-shocked individuals and Samples 1C and 3C were control individuals.

| Sample | Raw Reads | Filtered Reads | Mapped Reads | Filtered Mapped Reads |
| --- | --- | --- | --- | --- |
| 3E | 25259714 | 23736708 | 12499756 | 5682642 |
| 4E | 24794376 | 23304626 | 12619770 | 5843708 |
| 1C | 22749260 | 21494102 | 12499756 | 4717835 |
| 3C | 25485364 | 23900618 | 12594813 | 5745840 |

**Supplemental Table 5.** Exome capture specificity with a 10X threshold. Sensitivity measured here is the percentage of targets with at least 10 reads mapping successfully. Here, targets are broken up into subsets: All annotated exons, exons with at least 20X coverage from the RNA library, exons with at least 35X coverage from the RNA library, and exons with at least 50X coverage from the RNA library. EC_2, EC_4, and EC_7 are the three replicate captures with the original probe pool, and EC_1, EC_3, and EC_12 are the replicate captures with the probe pool exposed to 12 extra rounds of PCR.

| 10X | Capture Pool |  |  |  |  |  |
| --- | --- | --- | --- | --- | --- | --- |
| Targets | EC_2 | EC_4 | EC_7 | EC_1 | EC_3 | EC_12 |
| All Exons | 58.8% | 51.9% | 52.2% | 52.3% | 58.7% | 54.2% |
| 20XR Exons | 98.2% | 96.7% | 97.1% | 96.6% | 98.2% | 97.6% |
| 35XR Exons | 99.1% | 98.6% | 98.8% | 98.6% | 99.0% | 98.9% |
| 50XR Exons | 99.3% | 99.0% | 99.1% | 99.0% | 99.2% | 99.2% |

**Supplemental Table 6.**
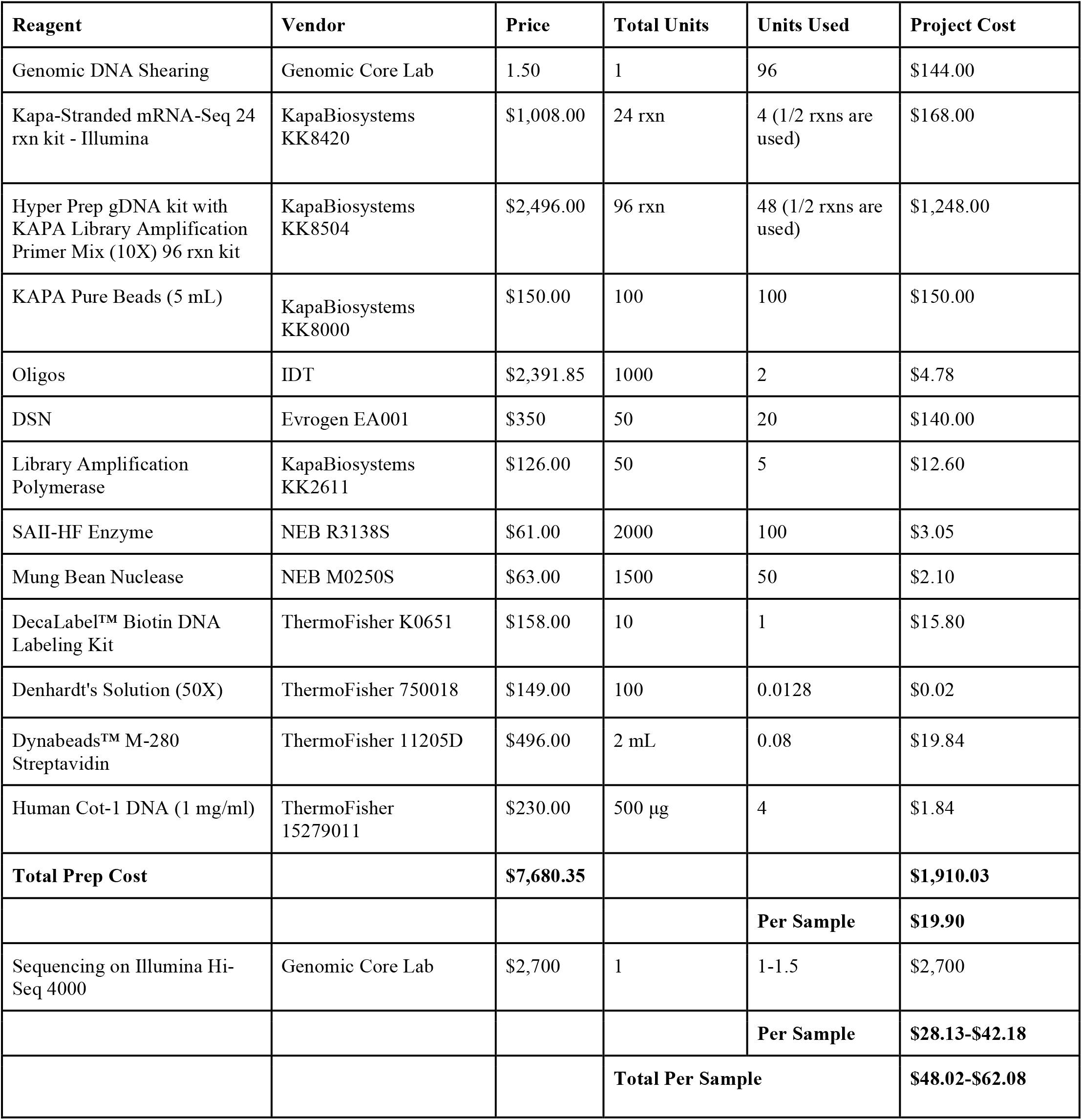
Per sample cost calculations for EecSeq. The calculations below assume 8 mRNA libraries are used to create probes to capture 96 samples in 8 capture reactions (12 samples per capture). Costs assumes the captured DNA is sequenced in one to one and a half lane(s) of Illumina High Seq 4000. This assumes that coverage levels for 96 samples in one lane would be equivalent to coverage levels seen in six pools of eight samples in half a lane. Cost does not include DNA or RNA extraction. See the github respository for more information on library preps and capture. Based on results in the main paper, this multiplexing strategy would give ~7.66x coverage per individual at exons represented by 35X sequencing depth at RNA-derived probes and would give ~3,508 exome SNPs at 10X coverage. Whether 96 individuals can be sequenced to enough depth in a single lane will depend on the number of megabases represented by the probes, the desired read depth, and the sensitivity and specificity of capture in the focal species. We have included the cost if 96 samples were sequenced over one and half lanes to include this potential variance.

## Supplemental Figures

**Supplemental Figure 1.**
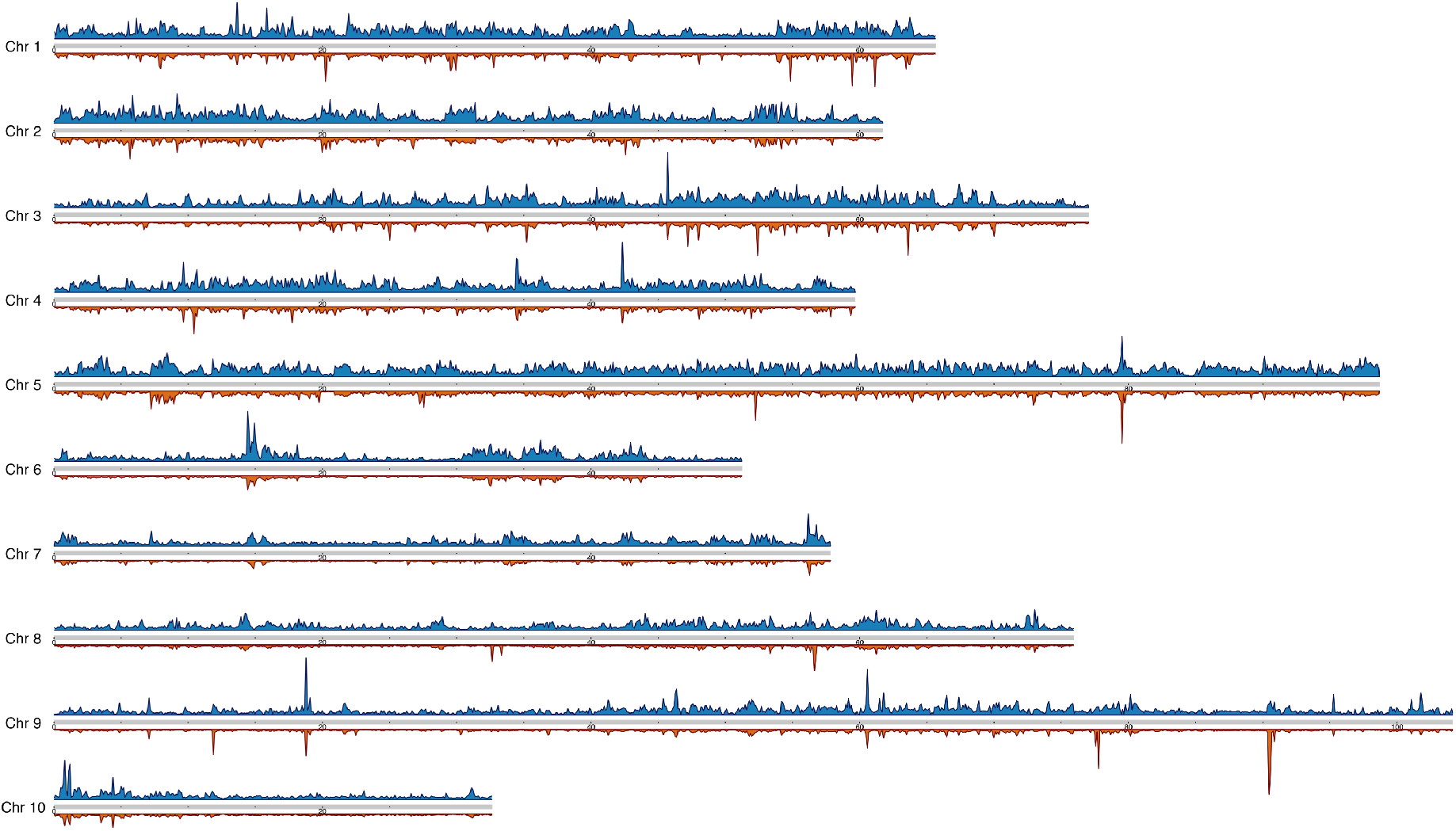
Expressed exome capture reads and RNAseq reads from probes plotted across the eastern oyster genome. Read coverage density was plotted in 10,000 bp sliding windows for both total RNA reads (red; below chromosome) and total EecSeq reads (blue; above chromosome) using the karyoploteR package (https://bioconductor.org/packages/release/bioc/html/karyoploteR.html).

**Supplemental Figure 2.**
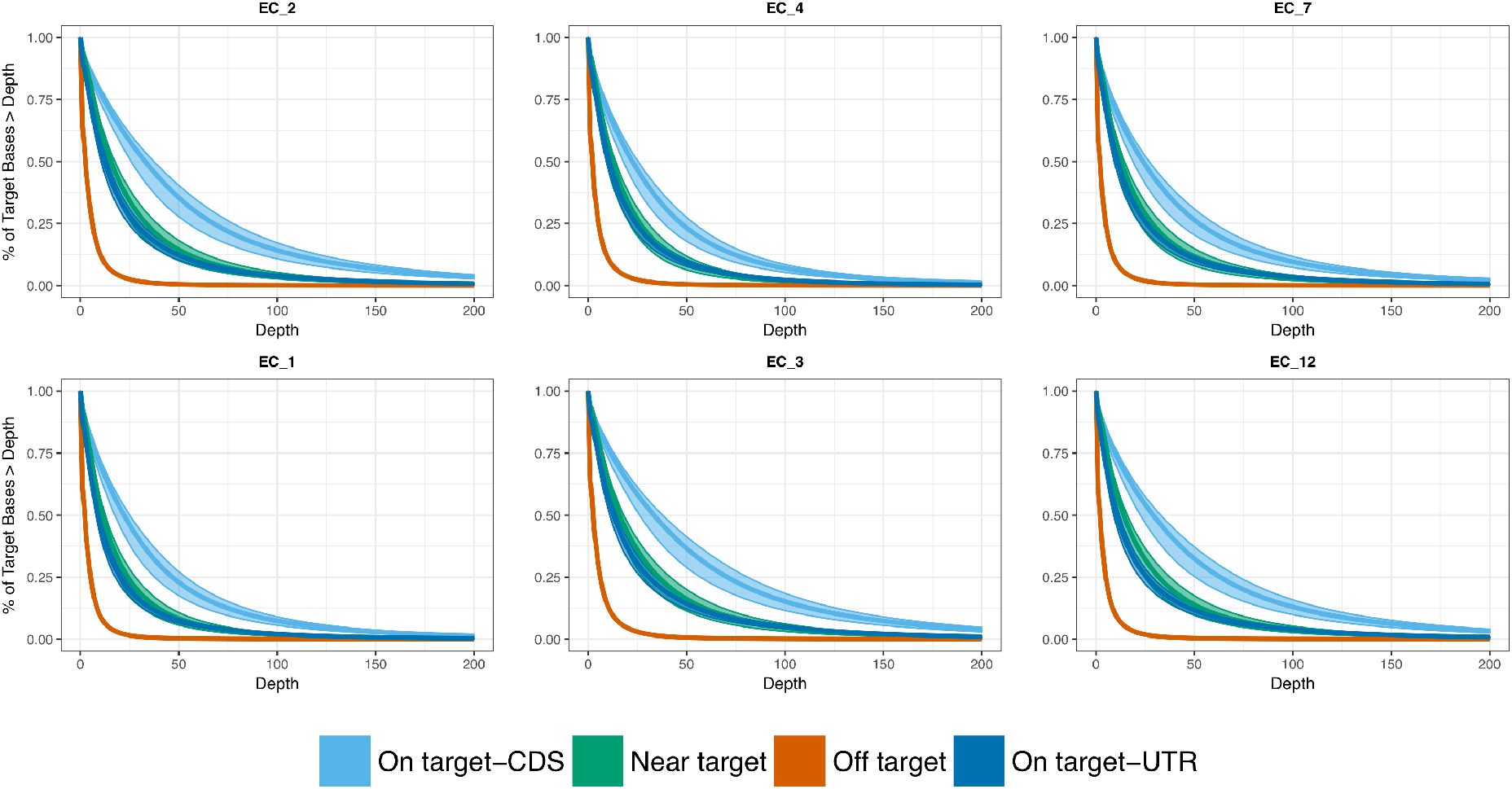
Per base pair sensitivity plot of EecSeq captures including CDS and UTR. To compare EecSeq to other capture methods, capture targets were defined as exons that had more than 35X coverage in the RNAseq (probe) data and confidence intervals were generated by defining capture targets between 20X RNAseq coverage and 50X RNAseq coverage. Near-target mapping were 150 bp on either side of the defined targets. For this figure, target exons were broken into coding sequence (CDS) and untranslated regions (UTR) for comparisons. This range corresponds to the modal DNA fragment length used for the capture libraries with the expectation that exon probes could capture reads that far from the original target. EC_2, EC_4, and EC_7 are the three replicate captures with the original probe pool, and EC_1, EC_3, and EC_12 are the replicate captures with the probe pool exposed to 12 extra rounds of PCR.

**Supplemental Figure 3.**
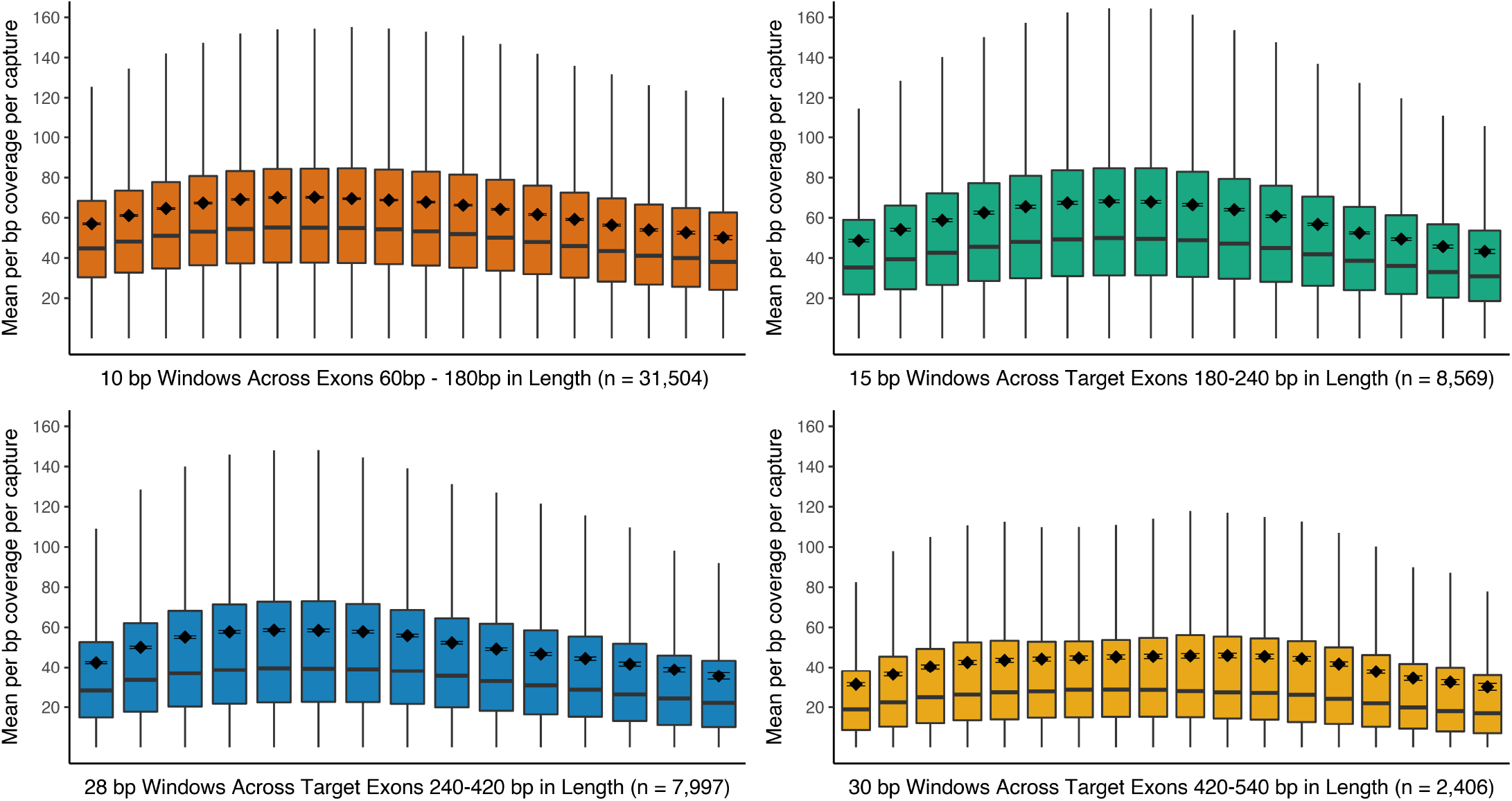
Boxplots of mean per basepair coverage levels plotted across target exons size windows. Target exons (those with at least 35X coverage in the RNA data) were broken into 10bp - 30 bp windows depending on overall size and the mean per basepair coverage per capture was calculated for each window size. The line each box represents the median of mean coverage values and the box surrounds the 25^th^ and 75^th^ percentiles. The mean of each bin class is plotted as a black diamond with standard error bars around it.

**Supplemental Figure 4.**
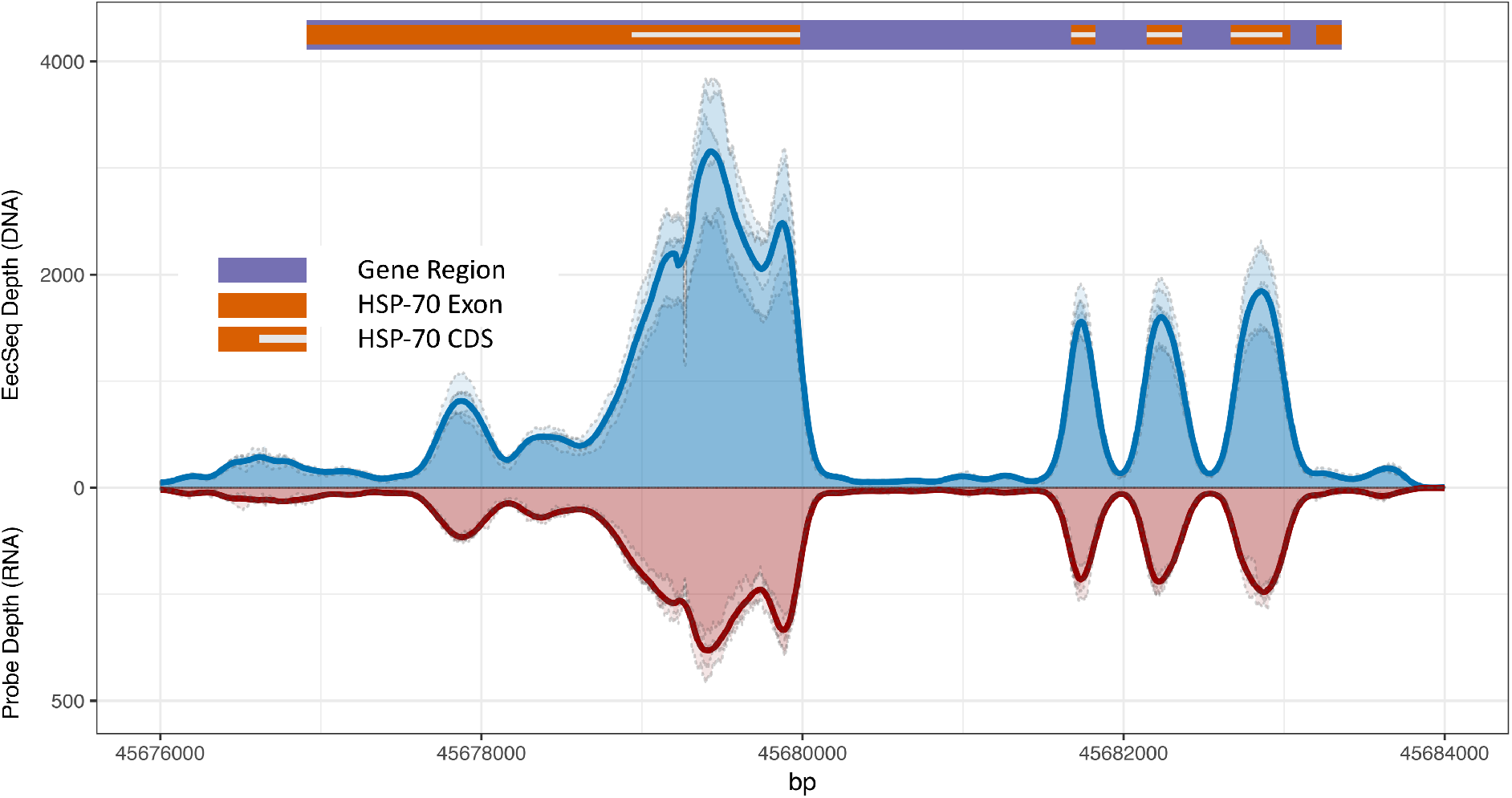
EecSeq capture and probe coverage across Heat Shock cognate 70 kDa. Coverage for each replicate capture pools is plotted along basepairs 45,760,000 to 45,684,000 of reference Chromosome NC_035782.1 containing the full gene region of 70 kDa protein 12B-like (NCBI Reference Sequence: XM_022468697.1). Each exome capture pool coverage is plotted in light blue with dashed grey border and a rolling 100 bp window average across all pools is plotted in dark blue. Each RNAseq (probe) sample coverage is plotted in light red with dashed grey border and a rolling 100 bp window average across all pools is plotted in dark red. Gene regions are marked in purple with exons color coded by gene. Coding sequence (CDS) is marked by a white bar within exon markers.

## References

Buerkle CA, Gompert Z (2013) Population genomics based on low coverage sequencing: how low should we go? Molecular Ecology, 22, 3028–3035.

Bi K, Vanderpool D, Singhal S et al. (2012) Transcriptome-based exon capture enables highly cost-effective comparative genomic data collection at moderate evolutionary scales. BMC genomics, 13, 403.

Bolger AM, Lohse M, Usadel B (2014) Trimmomatic: a flexible trimmer for Illumina sequence data. Bioinformatics, 30, 2114–2120.

Catchen JM, Hohenlohe PA, Bernatchez L et al. (2017) Unbroken: RADseq remains a powerful tool for understanding the genetics of adaptation in natural populations. Molecular ecology resources, 17, 362–365.

Chilamakuri CSR, Lorenz S, Madoui M-A et al. (2014) Performance comparison of four exome capture systems for deep sequencing. BMC genomics, 15, 449.

Christmas MJ, Biffin E, Breed MF, Lowe AJ (2017) Targeted capture to assess neutral genomic variation in the narrow-leaf hopbush across a continental biodiversity refugium. Scientific reports, 7, 41367.

De Mita S, Thuillet A-C, Gay L et al. (2013) Detecting selection along environmental gradients: analysis of eight methods and their effectiveness for outbreeding and selfing populations. Molecular ecology, 22, 1383–1399.

De Wit P, Pespeni MH, Palumbi SR (2015) SNP genotyping and population genomics from expressed sequences - current advances and future possibilities. Molecular ecology, 24, 2310–2323.

Dobin A, Davis CA, Schlesinger F et al. (2013) STAR: ultrafast universal RNA-seq aligner. Bioinformatics, 29, 15–21.

Garrison E, & Marth G (2012) Haplotype-based variant detection from short-read sequencing. arXiv, arXiv: 1207.3907

Fariello MI, Boitard S, Naya H, SanCristobal M, Servin B (2013) Detecting signatures of selection through haplotype differentiation among hierarchically structured populations. Genetics, 193, 929–941.

Hebert FO, Renaut S, Bernatchez, L (2013) Targeted sequence capture and resequencing implies a predominant role of regulatory regions in the divergence of a sympatric lake whitefish species pair *(Coregonus clupeaformis)*. Molecular Ecology, 22: 4896–4914.

Hoban S, Kelley JL, Lotterhos KE et al. (2016) Finding the Genomic Basis of Local Adaptation: Pitfalls, Practical Solutions, and Future Directions. The American naturalist, 188, 379–397.

Jones MR, Good JM (2016) Targeted capture in evolutionary and ecological genomics. Molecular ecology, 25, 185–202.

Jones FC, Grabherr MG, Chan YF et al. (2012) The genomic basis of adaptive evolution in threespine sticklebacks. Nature, 484, 55–61.

Keinath MC, Timoshevskiy VA, Timoshevskaya NY et al. (2015) Initial characterization of the large genome of the salamander Ambystoma mexicanum using shotgun and laser capture chromosome sequencing. Scientific reports, 5, 16413.

Lee CE, Remfert JL, Opgenorth T et al. (2017) Evolutionary responses to crude oil from the Deepwater Horizon oil spill by the copepod Eurytemora affinis. Evolutionary applications, 10, 813–828.

Lepais O, Weir JT (2014) SimRAD: an R package for simulation-based prediction of the number of loci expected in RADseq and similar genotyping by sequencing approaches. Molecular ecology resources, 14, 1314–1321.

Li H, Durbin R (2009) Fast and accurate short read alignment with Burrows-Wheeler transform. Bioinformatics, 25, 1754–1760.

Li H, Handsaker B, Wysoker A et al. (2009) The Sequence Alignment/Map format and SAMtools. Bioinformatics, 25, 2078–2079.

Lotterhos KE, Whitlock MC (2015) The relative power of genome scans to detect local adaptation depends on sampling design and statistical method. Molecular ecology, 24, 1031–1046.

Lowry DB, Hoban S, Kelley JL et al. (2016) Breaking RAD: An evaluation of the utility of restriction site associated DNA sequencing for genome scans of adaptation. Molecular ecology resources.

Lowry DB, Hoban S, Kelley JL et al. (2017) Responsible RAD: Striving for best practices in population genomic studies of adaptation. Molecular ecology resources, 17, 366–369.

McCartney-Melstad E, Mount GG, Bradley Shaffer H (2016) Exon capture optimization in amphibians with large genomes. Molecular ecology resources, 16, 1084–1094.

McKinney GJ, Larson WA, Seeb LW, Seeb JE (2017) RADseq provides unprecedented insights into molecular ecology and evolutionary genetics: comment on Breaking RAD by Lowry et al. (2016). Molecular ecology resources, 17, 356–361.

Müller T, Freund F, Wildhagen H, Schmid KJ (2014) Targeted re-sequencing of five Douglas-fir provenances reveals population structure and putative target genes of positive selection. Tree genetics & genomes, 11.

Nadeau NJ, Whibley A, Jones RT et al. (2012) Genomic islands of divergence in hybridizing Heliconius butterflies identified by large-scale targeted sequencing. Philosophical transactions of the Royal Society of London. Series B, Biological sciences, 367, 343–353.

Neves LG, Davis JM, Barbazuk WB, Kirst M (2013) Whole-exome targeted sequencing of the uncharacterized pine genome. The Plant journal: for cell and molecular biology, 75, 146–156.

Pastinen T (2010) Genome-wide allele-specific analysis: insights into regulatory variation. Nature reviews. Genetics, 11, 533–538.

Portik DM, Smith LL, Bi K (2016) An evaluation of transcriptome-based exon capture for frog phylogenomics across multiple scales of divergence (Class: Amphibia, Order: Anura). Molecular ecology resources, 16, 1069–1083.

Puritz JB, Matz MV, Toonen RJ et al. (2014), Demystifying the RAD fad. Molecular ecology, 23: 5937–5942.

Puritz JB, Hollenbeck CM, Gold JR (2014) dDocent: a RADseq, variant-calling pipeline designed for population genomics of non-model organisms. PeerJ, 2, e431.

Quinlan AR (2014) BEDTools: The Swiss-Army Tool for Genome Feature Analysis. Current protocols in bioinformatics /editoral board, Andreas D. Baxevanis…[et al.], 47, 11.12.1–34.

Reid NM, Proestou DA, Clark BW et al. (2016) The genomic landscape of rapid repeated evolutionary adaptation to toxic pollution in wild fish. Science, 354, 1305–1308.

Samuels DC, Han L, Li J et al. (2013) Finding the lost treasures in exome sequencing data. Trends in genetics: TIG, 29, 593–599.

Schlötterer C, Tobler R, Kofler R, Nolte V (2014) Sequencing pools of individuals — mining genome-wide polymorphism data without big funding. Nature reviews. Genetics, 15, 749–763.

Schmid S, Genevest R, Gobet E et al. (2017) HyRAD-X, a versatile method combining exome capture and RAD sequencing to extract genomic information from ancient DNA. Methods in ecology and evolution / British Ecological Society.

Suchan T, Pitteloud C, Gerasimova NS et al. (2016) Hybridization Capture Using RAD Probes (hyRAD), a New Tool for Performing Genomic Analyses on Collection Specimens. PloS one, 11, e0151651.

Sulonen A-M, Ellonen P, Almusa H et al. (2011) Comparison of solution-based exome capture methods for next generation sequencing. Genome biology, 12, R94.

Suren H, Hodgins KA, Yeaman S et al. (2016) Exome capture from the spruce and pine giga-genomes. Molecular ecology resources, 16, 1136–1146.

Syring JV, Tennessen JA, Jennings TN et al. (2016). Frontiers in plant science, 7, 484.

Therkildsen NO, Palumbi SR (2017) Practical low-coverage genomewide sequencing of hundreds of individually barcoded samples for population and evolutionary genomics in nonmodel species. Molecular ecology resources, 17, 194–208.

Yeaman S, Hodgins KA, Lotterhos KE et al. (2016) Convergent local adaptation to climate in distantly related conifers. Science, 353, 1431–1433.

Zhou L, Holliday JA (2012) Targeted enrichment of the black cottonwood (Populus trichocarpa) gene space using sequence capture. BMC genomics, 13, 703.

